# Docosahexaenoic acid (DHA), a nutritional supplement, modulates steroid insensitivity in asthma

**DOI:** 10.1101/2020.07.01.181354

**Authors:** Lipsa Panda, Atish P Gheware, Ashish Jaiswal, Dhurjhoti Saha, Bapu Koundinya Desiraju, Rakhshinda Rehman, Archita Ray, Joytri Dutta, Sabita Singh, Manish Kumar Yadav, Divya Tej Sowpati, Samit Chattopadhyay, Madhunapantula V. SubbaRao, Padukudru Anand Mahesh, Y. S. Prakash, Shantanu Chowdhury, Anurag Agrawal, Balaram Ghosh, Ulaganathan Mabalirajan

## Abstract

Asthmatics with poor steroid responsiveness are now found to use health services at higher frequency and contribute to socio-economic burden disproportionately. We have previously shown that a ω-6 fatty acid metabolite leads to a severe and steroid insensitive asthma-like condition in mice. Here, we investigated the role of retinoid-x-receptor gamma (RXRγ) and Docosahexaenoic acid (DHA), a ω3 fatty acid rexinoid ligand of RXR, on the features of steroid insensitivity in asthmatic condition. RXRγ was found to be reduced in the lungs of human asthmatics and mice with steroid insensitive allergic airway inflammation. RXRγ knockdown in naïve mice led to spontaneous asthma like features whereas RXRγ knockdown in allergic mice led to steroid insensitive asthma features. We observed while RXRγ binds to the glucocorticoid receptor (GR) gene and regulates its transcription, DHA increases the GRα expression in human bronchial epithelial cells and reverses the steroid insensitive features in mice with allergic airway inflammation. Docosahexaenoic acid (DHA), a ligand of RXR, was reduced in the sera of steroid-insensitive asthmatics. We conclude that DHA may prove to be a promising steroid sensitizing agent for the treatment of steroid insensitive asthmatics.

**Summary:** The molecular regulation of glucocorticoid receptor by retinoid-x-receptor gamma (RXRgama) has an implication in steroid insensitive asthma as we found that Docosahexaenoic acid (DHA), a nutritional supplement and natural ligand of RXRgamma, improves steroid sensitivity in steroid insensitive mice model of asthma and DHA levels are found to be low in steroid insensitive asthmatic patients.

## Introduction

Steroid resistance is one of the major menaces during the treatment of severe inflammatory diseases including asthma. Although only 5-10% of asthmatics do not respond to steroids, these asthmatics are responsible for the consumption of significant health resources and also for substantial morbidity and mortality (1). While the complete resistance to steroids is a rare entity, poor response to steroids is observed usually, where a high dose of steroids is required to control asthma – a condition termed as steroid dependent asthma (1). In addition to this, side effects caused by excessive use of steroids dramatically impact the quality of life.

Asthma, a disease which was primarily thought to be a Th2 dominant disorder, is now recognized as a syndrome with multiple phenotypes and heterogeneity (2). Though it has multiple phenotypes, it can also be classified into two larger phenotypes: high Th2 and low Th2 asthma. While the former phenotype predominantly embodies eosinophilic inflammation, the latter phenotype is not studied well enough (3, 4). Low Th2 phenotype asthmatics, such as very late onset asthmatics, obesity-associated asthmatics as well as asthmatic smokers and neutrophilic asthmatics show little allergic inflammation and respond poorly or do not respond to steroids (5, 6). Also, with the emergence of obese-asthma and non-eosinophilic asthma phenotypes, which are majorly refractory to steroid treatment, there is an imperative need for understanding the mechanisms of steroid resistance that would be an overarching phenomenon for phenotype independent steroid resistant/insensitive asthma (7). So far, various mechanisms have been demonstrated for the defective function of the steroids, like decreased levels of GR expression, reduced binding affinity of steroid with GR, decreased nuclear translocation, decreased capability of GR binding with GR elements, and participation of other transcription factors like NF-κB. In this context, several mediators were found to be involved in steroid resistance such as p38 mitogen-activated protein kinase (p38-MAPK), c-Jun N-terminal kinase 1, extracellular signal regulated kinase, histone deacetylase, etc (1, 8).

Although certain downstream activators of IL-13 like p38-MAPK are found to be involved in steroid resistance, the role of 15-LOX, an enzyme that oxygenates polyunsaturated fatty acids (also inducible by IL-13 through STAT-6) and its metabolites were not studied in relation to severe or steroid insensitive asthma. *13*-hydroxyoctadecadienoic acid (13-S-HODE), a 15-lipoxygenase metabolite of ω-6 fatty acid (linoleic acid), had been shown to cause spontaneous asthma like features in guinea pig and mice (9). In addition, the increase of 13-S-HODE in BAL fluids, sputum and lungs of asthmatic patients is well known (9, 10). In this context, our lab has shown recently that the administration of 13-S-HODE to mice with allergic airway inflammation (AAI) led to steroid insensitive asthma like features with neutrophilia (11). It was also observed that HODE treated human bronchial epithelia had reduced GRα transcript levels and activity. Further, NF-κB was observed to be partially involved in HODE mediated steroid resistance in mice (11). In the present study, we investigated the plausible mechanism of steroid resistance using HODE treatment in a mice model of asthma. We show that RXRγ is the only differentially expressed nuclear receptor gene in the lungs of mice with steroid insensitive asthma-like features and it can further modulate steroid sensitivity in mice. We also provide evidence that RXRγ can bind to intron 1 of the GR gene and positively regulates its expression, indicating it to be a plausible pathway for modulating steroid sensitivity in humans. In this respect, retinoids and steroids have been shown to converge at various molecular pathways (12– 13). Rexinoid receptors or retinoid X receptors, a member of class II nuclear receptor family is considered to be the master regulator due to its role in cellular and organ differentiation, apoptosis and various metabolic processes (14). Their activities are known to be controlled by their expression levels and activation by their endogenous ligands such as 9-cis retinoic acid and docosahexaenoic acid (DHA, ω-3 fatty acid) (15). In this study, we show for the first time, that DHA was reduced in human sera of steroid insensitive asthmatic patients as compared to steroid-sensitive patients and it also modulates steroid sensitivity in mice with allergic airway inflammation.

## Methods

### Study design

The goal of this study was to determine the effects of RXR gamma and its ligand, DHA, in steroid insensitive asthma like features in mice. Based on the observations of reduced expression of RXRγ in steroid insensitive mice lung and RXR/RAR binding sites in glucocorticoid receptor gene, we hypothesized that RXRγ reduction in asthmatic lungs may drive glucocorticoid insensitivity. Using human lung biopsies, in vitro studies using human bronchial epithelia, and mice models, we explored the effect of RXR gamma and its ligand, DHA in steroid insensitive asthma like features in mice.

### Mice studies

All mentioned animal protocols were approved formally by the Institutional Animal Ethical Committee at the Institute of Genomics and Integrative Biology (IGIB) and Indian Institute of Chemical Biology (IICB). The male BALB/c mice (6-8 weeks) were procured from Central Drug Research Institute, Lucknow, India and IICB, Kolkata, India and maintained in the individual ventilated cages in IGIB, Delhi, India, and IICB. All mice experiments were performed in accordance with the relevant guidelines and regulations of the Committee for the Purpose of Control and Supervision of Experiments on Animals (CPCSEA).

### Mice models

Seven different models were utilized as mentioned in Supplementary Table 1: I) HODE induced steroid insensitive allergic mice model, II) Cockroach allergen extract (CE) induced airway inflammation model, III) RXRγ knockdown model in naïve mice, IV) RXRγ knockdown induced steroid insensitive allergic mice model using ovalbumin allergen, V) RXRγ knockdown induced steroid insensitive allergic mice model using cockroach allergen, VI) Effects of DHA/ATRA in HODE induced steroid insensitive allergic mice model, and VII) Effects of DHA in RXRγ knockdown induced allergic mice model. The details of each model have been explained in respective results section and also in Supplementary Table 1.

### Allergen-immunization and challenge

For ovalbumin induced allergic airway inflammation models (I, IV, VI and VII), mice were sensitized with three intraperitoneal injections of 50µg OVA adsorbed in alum for three weeks and challenged with 3% OVA in PBS for 7 days as described earlier (9, 16, 17). For cockroach allergen extract (CE) induced allergic airway inflammation models (II and V), naïve mice were exposed to PBS or cockroach allergen [different concentrations (10µg, 50µg, and 100µg) for mice model II and 50µg for mice model V] as described earlier (18). After brief isoflurane anaesthesia, PBS or cockroach allergen extract dissolved in PBS was given intranasally on Days 1-5. After 5 days, PBS or cockroach allergen extract was given intranasally in Day 11-14 followed by AHR measurement, BAL procedure and euthanasia on Day 15. The whole body cockroach (*Periplaneta Americana*) allergen extracts were obtained from Greer Laboratories as endotoxin free lyophilized extracts and also as a gift from Dr Naveen Arora, CSIR-IGIB, Delhi, who had isolated Per a 10 from cockroach extracts (19), demonstrated the role of cockroach allergen in causing airway epithelial cell activation (20) and also had demonstrated the beneficial effects of protease inhibitor in reducing cockroach allergen mediated allergic airway inflammation (21).

### Administration of 13-S-HODE, Dexamethasone, RXRγ siRNA, RXRγ plasmid, DHA, and ATRA

13-S-HODE (Cayman, Michigan, USA) or VEH (50% ethanol) was instilled into the nasal openings of each isoflurane anesthetized mouse (Models I and VI). Based on our previous publication (11) we have selected the dose of 0.6mg/kg for each mouse. 13-S-HODE was administered intranasally on days 24, 26 and 28 as shown in Figure S5A. Dexamethasone (Sigma-Aldrich, MO, USA), dissolved in 50% ethanol, was given orally (0.75mg/kg) to mice from day 24 to 28 as shown in Figure S5A. RXRγ siRNA (25μg-100μg) or scrambled siRNA was instilled to the nasal openings of each isoflurane anesthetized mouse as shown in Figure S2A (Days 1 & 3), Figure S5D (Days 24 and 26) in Models III, IV and VII and Days 11 & 13 in case of CE Model 5. RXRγ plasmid (25μg) or VEH (empty plasmid) was given intravenously to each mouse on days 1 and 3 in Model II (Figure S2A). ATRA (dissolved in DMSO, 10mg/kg), DHA (dissolved in corn oil, 12.5mg/kg) or DMSO/corn oil were administered orally and intraperitoneally, respectively, from day 24 to 28, 2hrs after the dexamethasone treatment as shown in Figures S5A, D. Each experiment had negative controls in which vehicle in which particular compound/metabolite was dissolved like 50 % ethanol (for 13-S-HODE or dexamethasone), scrambled siRNA (for RXRγ siRNA), control empty plasmid (for RXRγ plasmid), DMSO (for ATRA) and corn oil (for DHA). In some experiments, we have used more than one vehicle depends on the type of experiment (Supplementary Table 1).

### Airway hyperresponsiveness measurement, bronchoalveolar lavage (BAL) and Lung histopathology

Airway hyper-responsiveness was measured with invasive flexiVent (SCIREQ, Montreal, Canada) on Day 5 (Model III), Day 15 (Models II & V) and Day 28 (Models I, IV, VI, & VII).

Bronchoalveolar lavage (BAL) was performed to separate cell pellets that were used to measure both differential and total cell counts (9, 16, 17). From these two cell counts, absolute cell counts were calculated by multiplying them (9, 16, 17). After BAL procedure, mice were euthanized using higher dose of thiopentone sodium and lungs were removed. Formalin-fixed lung sections were stained with Haematoxylin & Eosin (H & E) and Periodic acid-Schiff stainings to determine the airway inflammation and goblet cell metaplasia, respectively (9, 16, 17). The total inflammation score was calculated as described earlier (16). Briefly, random numbered H & E stained lung slides were evaluated by experimentally blind investigators. Scores, 0-4, were given for both perivascular and peribronchial inflammation, Score 0: No detectable inflammation; Score 1: occasional cuffing of bronchi or vessels with inflammatory cells; Score 2: when bronchi or vessels were cuffed with 1–3 layers of inflammatory cells; Score 3: when bronchi or vessels were cuffed with 3–5 layers of inflammatory cells; and Score 4: when bronchi or vessels were cuffed with more than 5 layers of inflammatory cells. Both perivascular and peribronchial scores were summed to get the total inflammation score.

### ELISAs

Mouse RXRγ (MyBioSource), Human RXRγ, DHA, retinol (Cloud Clone Corp), KC, IL-13, IL-17A, IL-21, IL-22 (E-Biosciences, CA, USA), IL-4, IL-5, IFNγ, and OVA specific IgE (BD Pharmingen) were performed according to the manufacturer’s instructions from human lung homogenates (RXRγ), human sera (DHA, retinol), mice sera (OVA specific IgE) and mice lung homogenates (remaining parameters). The optimization has been done using different concentrations of mice lung homogenates [5µg (IL-13 & KC), 7.5µg (IL-17), 10µg (IL-5), 20µg (IL-4), 20µg (IFNγ)] and so the results are expressed in pg/5-40µg Lung.

For measuring GR activity (Active Motif, CA, USA), BEAS-2B cells obtained after HODE and DEX treatment (as described above), were processed for nuclear extract. Elisa was performed using manufacturer’s protocol with 20μg of nuclear extract. The measured optical density is plotted in arbitrary units (AU) in fold change, calculated by optical density test/optical density veh.

### Human subjects

Lung samples and sera samples from human asthma patients were obtained from St. Marys Hospital at Mayo Clinic Rochester and JSS Medical College, Mysore, India, respectively. The post-surgical lung samples were collected from 50-75 years old patients after surgical indications about various etiologies like focal nodules of benign or cancer (Supplementary Table 2). The clinical examinations, lung function with bronchodilator responses were performed to diagnose asthma and mild-moderate cases were selected for the study. There were 42 sera samples (25 healthy controls, 8 steroid sensitive and 9 steroid insensitive). The mean age for steroid sensitive and steroid insensitive was 45 ± 14.62 and 41 ± 6.72 yrs, respectively. The duration of steroid dose for steroid sensitive and steroid insensitive was 6 ± 6.422 and 63.6 ± 56.27 months, respectively. Ethical committee approval from the Review Boards of respective hospitals were obtained and also from IGIB to use these samples.

### In vitro experiments

Human bronchial epithelial cells (BEAS-2B) were obtained (ATCC, Manassas, VA, USA) and maintained in HAM’s F12 (Sigma-Aldrich, MO, USA) with 10% fetal bovine serum. Human breast adenocarcinoma cell line (MCF-7) were obtained (ATCC, Manassas, VA, USA) and maintained in DMEM-F12 (Sigma-Aldrich, MO, USA) with 10% fetal bovine serum. The cells were pretreated with dexamethasone (10^−6^ M, Sigma, MO, USA) for 3hrs before stimulating with vehicle (ethanol) or 13-S-HODE (35 μM, Cayman, Ann Arbor, Michigan, USA) for 16 hrs. These cells were harvested for further experiments and supernatants were stored for cytokine assays.

For transfection assays, 1 million cells were seeded on 6 well plates. RXRγ siRNA (sc-270445, Santacruz) 150nM, was transfected into cells using lipofectamine 2000 in OPTIMEM media. Media was changed with Hams-F12 after 5 hours, and cells were cultured for 48hrs. Dexamethasone (10^−6^M) was added to the fresh media after 5hrs of transfection followed by addition at regular intervals of 24 hrs. Later, cells were harvested and processed for either western analysis, real-time analysis.

### Quantitative Real-Time PCR

Beas-2B cells were cultured in Hams-F12 with 10% FBS with or without 10^−6^M agonist (ATRA, DHA, 9CRA)/vehicle (DMSO) for 48 hours. These cells along with cells treated with RXRγ siRNA and DEX (as mentioned in cell culture) were subjected to RNA isolation, performed by RNeasy mini kit, (Qiagen). cDNA synthesis was carried out by EZ first strand cDNA synthesis kit (Applied Biosciences). Real time PCR was run using Kapa SYBR FAST and carried out in the Roche LC480 real time machine. Following are the primers used for transcript analysis: huRXRG : **FP**-5’GGGAAGCTGTGCAAGAAGAAA3’ **RP**-5’ TGGTAGCACATTCTGCCTCACT 3’, hu GRA : **FP**-5’ ACGGTCTGAAGAGCCAAGAG3’, **RP**-5’ CAGCTAACATCTCGGGGAAT 3’, huGRB: **FP** 5’ACGGTCTGAAGAGCCAAGAG 3’, **RP**-5’TGTGTGAGATGTGCTTTCTGG3’; huRXRA : **FP**-5’CGACCCTGTCACCAACATTTGC3’, **RP**-5’GAGCAGCTCATTCCAGCCTGCC3’; huRXRB-**FP**: 5’ATACTCTTGCCGGGACAACA3’, **RP** : 5’TCCCCATCCTTGTCCTTTCC3’; b-Actin: **FP**-5’CCAACCGCGAGAAGATGA3’, **RP**-5’CCAGAGGCGTACAGGGATAG3’.

### Quantitative Real-Time PCR array

Nuclear receptor array, RT^2^ Profiler PCR Array, (Qiagen) was performed according to manufacturer’s protocol in the total lung mRNA of HODE-treated steroid resistant mice. To do so, we performed a nuclear receptor PCR array which comprised of 86 genes with the total mRNA isolated from the HODE induced steroid resistant mouse lungs. We normalized the gene expression observed in groups OVA, OVA+HODE, OVA+HODE+DEX and OVA+DEX by OVA alone, to nullify the effect observed due to OVA.

### Immunoblotting and Immunohistochemistry

The cells were processed (mentioned in cell culture methods) and lysates were prepared to run sodium dodecyl sulfate–polyacrylamide gel electrophoresis. Following a transfer, membranes were probed with specific primary antibodies and corresponding secondary antibodies. After blocking the membranes with 5% BSA (sigma) for one hour, these were subjected to primary antibody incubation. Membranes were incubated in 1X PBST with 1:200 dilution of anti-RXRγ (abcam-15518), 1:200 dilution of anti-GRα (abcam-3579), 1: 400 dilution of RXRγ (sc-555x) and 1: 2000 dilution of GAPDH (sc-32233). The detection was done by colorimetric assay using DAB (3,3-Diaminobenzidine tetrahydrochloride) as a substrate. The expressions of RXRγ in lung sections were performed by immunohistochemistry as described earlier (11).

### Chromatin Immunoprecipitation

ChIP assays were performed as per protocol provided by Upstate Biotechnology with modifications as suggested in Fast ChIP protocol (6). 1 million of Beas-2B cells were used for the experiment and anti-RXRγ (abcam-15518, 10μg), anti-RAR (sc-773x, 10μg) were used to immunoprecipitate chromatin. Human IgG (sc-2907, 10μg) was used for mock immunoprecipitation in the cell line. Briefly, cells were fixed with 1% formaldehyde for 10 minutes. Chromatin was sheared to an average size of ∼500 bp using a Misonix 3000 sonicator (pulse 30 sec on and 45 sec off) for 30 mins. Twenty-five per cent of lysate was used to isolate input chromatin using phenol–chloroform and ethanol precipitation. Lysate was precleared using protein-A sepharose beads, and ChIP was performed using 10 μg of the respective antibody incubated overnight at 4°C. Immune complexes were collected using herring sperm DNA-saturated protein-A Sepharose and washed extensively. Chelex-100 resin was used to extract DNA from immunoprecipitated chromatin as described previously. Real time was performed to check the occupancy of the RXR’s and RAR with the following primers. RXRE Intron1: **FP**-5’AAGGCTGATGTTGTGTGCTG3’, **RP**: 5’TCCCTGCTGTATCTTCATTGC3’; Exon 1: **FP**-5’ GGAAATTGCAACACGCTTCT 3’, **RP**: 5’TATTCCTTCCCCACTCATGC3’; Negative control: **FP**-5’ CAAATCAGCCTTTCCTCGGG3’, **RP**-5’CTGGCCCTTCAAATGTTGCT3’; RXRE intron 2: **FP**: 5’AATTTGTAGGCCAGGCACAG3’, **RP**: 5’CCACCACACCTGGCTAATTT3’; PIAS3: **FP**-5’TGGCGGGACTCTGGGATTTC3’, **RP**-5’TAACGACGAGAAGGCGGACC3’. Fold enrichment was calculated by normalizing the test ct value with IgG ct value.

### Oligo-pull down assay

Oligonucleotide pull down assay was used (5) to probe direct binding of RXRγ with biotinylated oligonucleotides from NR3C1 Intron1. For pull down assays, 5μg of biotinylated wild type RXRE Binding site and as well as its mutant motif were incubated with 400μg of BEAS-2B nuclear extract for 1 hour at room temperature in binding buffer (15% Glycerol, 12mM HEPES, 4mM Tris, 150mM KCl, 1mM EDTA and 1mM DTT). The whole complex was incubated with streptavidin-agarose (Thermo Fisher catalogue no. 20349) for overnight at 4°C. In all experiments, the beads were washed three times with washing buffer (50 mM Tris-HCl [pH 7.5] and 150 mM NaCl) and the bound proteins were eluted by boiling in 5× sodium dodecyl sulfate (SDS) sample buffer (20 mM Tris-HCl [pH 6.8], 10% glycerol, 4% SDS, 100 mM dithiothreitol, 4 mM EDTA, and 0.025% Coomassie brilliant blue R250) and were subjected to western blot analysis using antibody from Abcam - ab15518 and sc-555x which recognizes RXRγ. The wild type sequence of RXRE oligonucleotide from NR3C1 promoter is the following: biotin-5’ FP-GAATTCAGGTCACTCAGAGGTCAAAGCTT; 5’ RP-AAGCTTTGACCTCTGAGTGACCTGAATTC the sequence which is underlined represents 17-mer RXRγ binding site. Mutant oligonucleotide sequence: biotin-5’FP – GAATTCAGCGACATGACGATCGAAAGCTT and 5’RP - AAGCTTTCGATCGTCATGTCGCTGAATTC; mutant/disrupted oligonucleotide served as binding control.

### Reporter gene assays and cell transfections

MCF-7 cells were transfected at 60-80% confluency with pEZX-PF02-01-eGFP (GR promoter) and pEZX-PF02-02-eGFP (17bp of RXRE deleted, GR (N) promoter) alone, or together with RXRγ siRNA (SC-270445, Santa Cruz, 150nM) or scrambled siRNA (Santa Cruz) using lipofectamine 2000 (Invitrogen) in DMEM-F12 media with 10% FBS. After 6 hrs, the medium was replaced with fresh medium and cells were incubated for 48hrs. These cells were then analyzed with flow cytometry for the geometric mean intensity of enhanced GFP.

For enhancer assays, MCF7 cells were transfected with CS-GS290L (HSVTK promoter(63bp)-GLuc, 0.5ug), pEZX-PG02-RXRE(MCS1-RXRE(153bp)-MCS2-HSVTK promoter(63bp)-GLuc, 0.5ug) and pEZX-PG02-SV40 (MCS1-SV40 enchancer (247bp)-MCS2-HSVTK promoter(63bp)-GLuc, 0.5ug) with pGL2-Firefly luciferase (Promega, 40ng) alone, or along with RXRγ siRNA (sc-270445, Santa Cruz,150nM) or scrambled siRNA (Santa Cruz) or/and RXRG plasmid (Myc-DDK-tagged, origene,NM_006917, 0.5ug) or pCMV6-Entry plasmid (Myc-DDK-tagged, origene, 0.5ug) using jetPRIME (Polyplus) in DMEM-F12 with 10% FBS. After 6hrs, the medium was replaced with fresh medium and agonists (ATRA, 9CRA and DHA)/ vehicle (DMSO) were added in 10^−6^M for 48hrs after transfection. Cells were lysed after the incubation was over and analysed for luciferase expression using Promega dual luciferase activity kit, wherein, Gaussia luciferase activity of each sample was normalized with firefly luciferase activity. Later, luciferase activity of RXRE-GLuc was normalized with TK-GLuc and plotted.

### Statistical analysis

All data represent mean ± SEM. The number of mice per each group and number of repeat experiments have been mentioned in Figure legends. *p <0.05, **p <0.01, ***p <0.001. A p-value more than 0.05 is considered non-significant (NS). Statistical significance of the differences between the two groups was determined with a two-tailed Student’s *t* test. For more clarification, the name of the statistical test for individual experiments has been mentioned in the respective figure legend. We have presented data of combined experiments or representative results of at least two independent experiments indicate that all the experiments were reproducible.

## Results

### RXRγ is decreased in the lungs of mice mimicking steroid insensitive asthma features and human asthmatic lungs

Earlier studies from our laboratory have demonstrated the steroid insensitive asthma features in the mice that were instilled intranasally with HODE (11). We have used the same steroid insensitive model (Model I mentioned in Supplementary Table 1) to explore the possible mechanisms underlying steroid resistance. There were five groups of mice as mentioned in Methods: SHAM, OVA, OVA+DEX, OVA+HODE and OVA+HODE+DEX. In our earlier study, we observed that OVA+HODE+DEX mice did not respond to steroid treatment compared to OVA+DEX (11) as the former group did not show any reduction in the airway hyperresponsiveness and airway inflammation.

The nuclear receptor array experiments revealed that Rxrγ and NF-κB2 were regulated differentially upon HODE administration. Rxrγ, but no other member of the RXR or RAR family, was found to be reduced in OVA+HODE+DEX mice when compared to OVA mice and was restored in OVA+DEX mice (Fig. S1A and Supplementary Table 3). However, NF-κB2 was increased in OVA+HODE+DEX mice as compared to OVA mice and was reduced in OVA+DEX mice (Supplementary Table 3). Prior studies from our laboratory have demonstrated elevated p-NF-kB in HODE induced steroid resistance in mice (11).

First, we validated the array experiments using immunohistochemistry. RXRγ indeed was found to be present modestly in bronchial epithelia of SHAM mice whereas its expression was found to be reduced in the bronchial epithelia of OVA induced asthmatic mice (Fig. 1A). In contrast, steroid (dexamethasone) treatment had restored the expression of RXRγ in bronchial epithelia (Fig. 1A1, A2). However, dexamethasone treatment could not restore the expression of RXRγ in bronchial epithelia of HODE administered mice. Notably, RXRγ was also expressed in sub-epithelial mesenchymal regions especially in HODE administered mice independent of the dexamethasone treatment.

**Figure 1.**
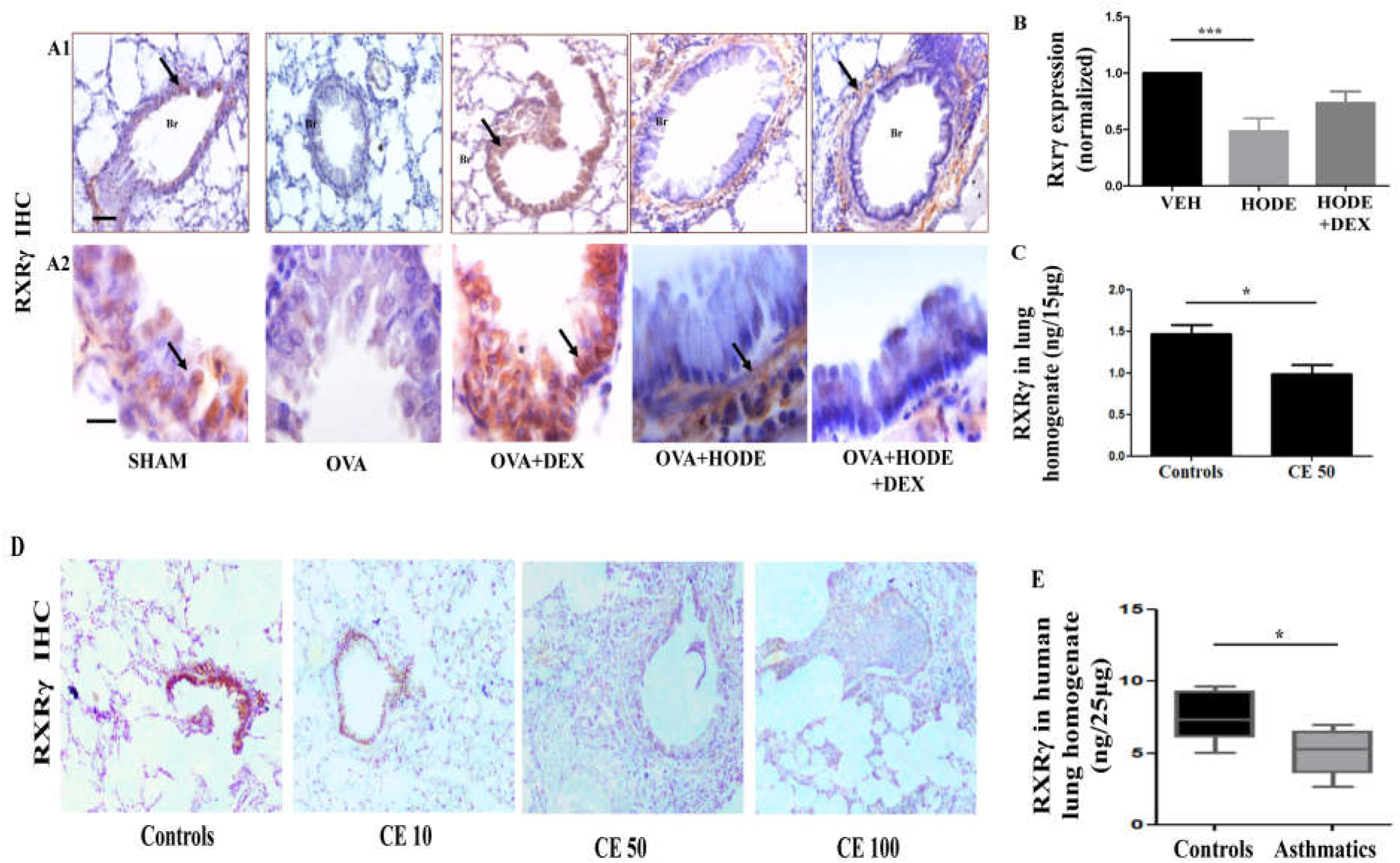
RXRγ is downregulated in lungs of human and asthmatics steroid insensitive asthmatic mouse. **A1 and A2)** Representative IHC images (20X magnifications for A1 and 100 X magnifications for A2) for the expression of RXRγ in HODE induced steroid resistance model (Model I). Bars=100µm (A1) and 25µm (A2). n=5-6 per each group and shown images are representative of two independent experiments. **B)** mRNA levels of RXRγ in Beas-2B cells treated with HODE (35uM) and DEX (10^−6^M) plotted in fold change with respect to housekeeping genes, beta actin and beta 2 microglobulin. This is combined data of 4 independent experiments and unpaired t test between VEH and HODE. **C**) Protein levels of RXRγ in lungs of CE induced allergic mice (unpaired t test, n=9 and this is combined data of 2 independent experiments). **D**) Representative IHC images (10X magnifications) for the expression of RXRγ in lungs of mice that have been induced with different concentrations of cockroach allergen extract (CE, Model II). n=4-6 per each group and shown images are representative of three independent experiments. **E)** Protein levels of RXRγ in asthmatic human lung homogenates (unpaired t test with Welch’s correction, n=8-9 per each group). Data represents mean ± SE; *p <0.05, ***p < 0. 001. Br: Bronchi. Arrows indicate the positive expression.

As RXRγ is expressed predominantly in bronchial epithelia of normal mice, we performed experiments in human bronchial epithelia. As shown in Fig. 1B, HODE induction alone was sufficient to reduce the expression of RXRγ in these cells whereas dexamethasone (DEX, 10^−6^M) was not able to restore RXRγ upon HODE induction.

To determine the clinical relevance of this finding, we examined the status of RXRγ protein in the lungs of cockroach allergen induced mice. There was also a reduction in the expression of RXRγ predominantly in bronchial epithelia in cockroach allergen extract (CE) induced mice with allergic airway inflammation (Fig. 1D, E). These CE exposed mice (with the different concentrations, 10 µg, 50 µg and 100 µg CE, Model II in Supplementary Table 1) had airway inflammation and goblet cell metaplasia along with an increase in AHR to higher doses of Methacholine like 50 mg/kg (Fig. S1, D-F).

Further, we also measured the levels of RXRγ human asthmatic lungs by ELISA. The levels of RXRγ were significantly lower in human asthmatic lungs compared to normal controls (Fig. 1C). These samples were from 50-75 years old patients who had undergone lung surgery for various etiologies like focal nodules of benign (Supplementary Table 2). Though, we have measured RXRγ in recently diagnosed asthmatic patients, we could not assess their steroid sensitive status. We will be performing the same in our future investigations. Also, there was no change in the expression of the other two isoforms of RXR, RXRα and RXRβ (Fig. S1B-C).

### RXRγ maintains lung homeostasis

Before examining the role of RXRγ in the steroid resistance, we determined its involvement in asthma pathogenesis as there is no available study to demonstrate such a physiological role. The RXRγ was found to be expressed in predominantly in bronchial epithelia of normal mice whereas it was found to be reduced in in asthmatic lungs. To test whether this reduction is a bystander effect or plays a causative role, we downregulated RXRγ in naïve mice (Model III, Supplementary Table 1) using different concentrations (25ug, 50ug, 75ug, and 100ug) of siRNA (Fig. S2A, B) and observed spontaneous airway hyper-responsiveness (AHR) in siRNA treated mice (Fig. S2F). These mice had shown airway inflammation even at 25μg dose (Fig. 2A, Fig. S2C). We have combined 25 and 50μg dose groups as one group, 25-50μg, since there was no difference in their effects. We observed an increase in the numbers of inflammatory cells like mononuclear immune cells such as monocytes & lymphocytes and neutrophils in BAL fluids of RXRγ siRNA administered mice (Fig. 2A, B). However, there was an insignificant increase in macrophages. While 25-50μg dose group had shown a significant increase in mononuclear immune cells, 75-100μg dose group had shown a significant increase in neutrophils compared to the scrambled siRNA group (Fig. 2B). The increase in AHR and airway inflammation in the RXRγ siRNA administered mice was associated with an increase in the levels of IL-17 family cytokines like IL-17, IL-21, and IL-22 (Fig. 2C). The 75-100μg dose group had shown a significant increase of these IL-17 family cytokines whereas 25-50μg dose group had a significant increase in IL-21 (Fig. 2C). Though the IL-13 levels were not significantly increased (Fig. 2C), IL-4 levels were found to be increased in RXRγ siRNA administered mice (Mean ± SEM; 17.3 ± 5.3pg/20µg lung in case of Scrambled siRNA 100 μg group and 73.9 ± 4.7 pg/20µg lung in 75-100μg RXRγ siRNA administered mice; p<0.001). On the other hand, the levels of IFNγ were found to be reduced significantly in 75-100μg RXRγ siRNA administered mice (Mean ± SEM; 478.1 ± 53.73 pg/40µg lung in case of Scrambled siRNA 100 μg group and 277 ± 48.61 pg/40µg lung in 75-100μg RXRγ siRNA group; p=0.0152). To confirm these results further and also to validate the specificity of RXRγ siRNA, we had upregulated RXRγ using RXRγ-plasmid in the RXRγ knockdown mice. We have performed pilot experiments with different concentrations (25 µg, 50 µg, and 75 µg) of RXRγ-plasmid in naïve mice (Fig. S2 D-E) and we selected 25 µg dose for further experiments. RXRγ-plasmid treatment reduced the features induced by RXRγ siRNA like spontaneous airway hyperresponsiveness to methacholine and airway inflammation (Fig. S2F-H).

**Figure 2.**
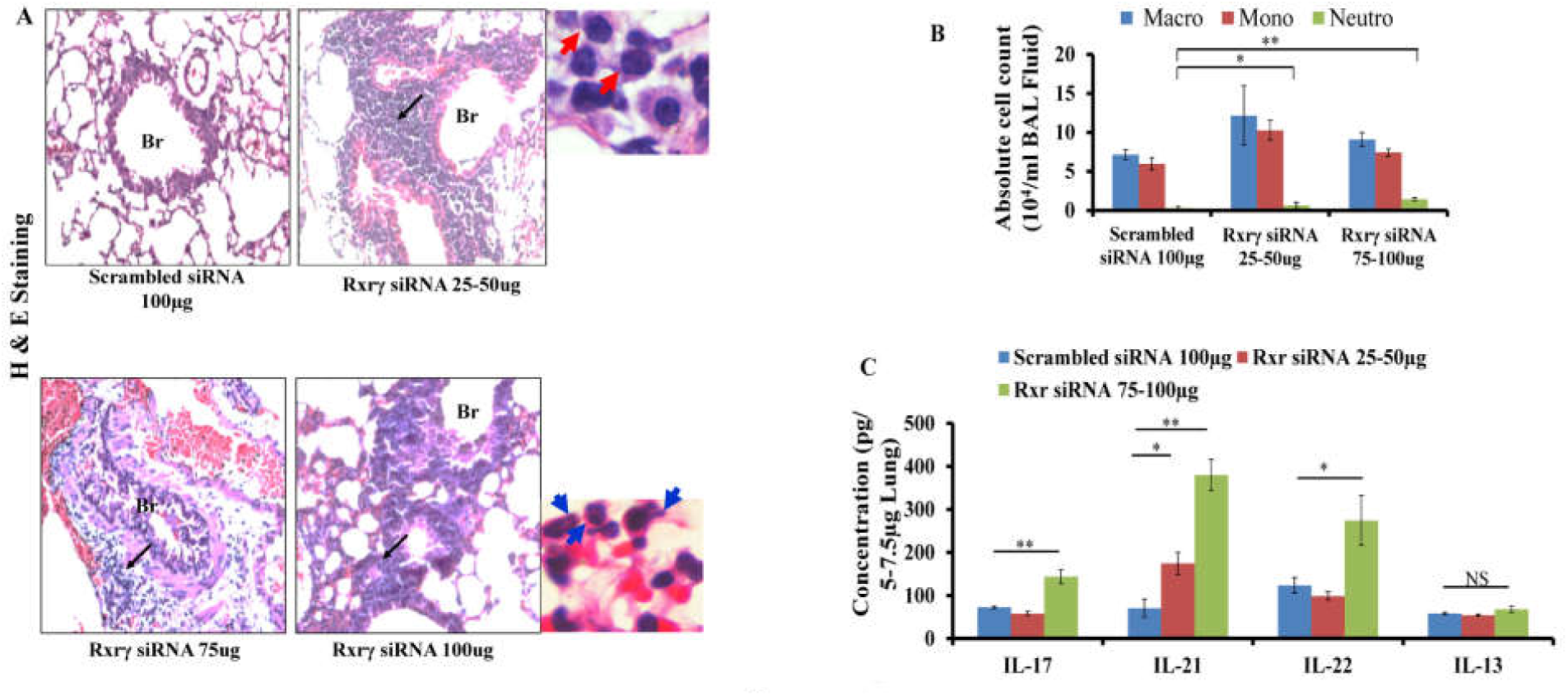
Knockdown of RXRγ impairs lung homeostasis and causes lung inflammation. **A**) Representative photomicrographs (20 X magnifications) of bronchovascular regions of different groups of mice stained with haematoxylin and eosin (H & E) in naïve mice downregulated with RXRγ siRNA. Black arrows indicate the inflammation in bronchovascular regions. Higher magnified images (100 X magnifications) near RXRγ siRNA 25-50 µg and RXRγ siRNA 100 µg images show the presence of mononuclear immune cells (red arrow) and neutrophils (blue arrow). Absolute cell count (**B**) in BAL fluids and the concentrations of various cytokines in the total lung lysates (**C**) (unpaired t test). Data represents mean ± SE; and shown data is representative two independent experiments and similar trend was observed in each independent experiment. *p <0.05, **p <0.01. Br: Bronchi.

### RXRγ knockdown leads to steroid insensitivity in allergic mice

#### Effect of RXRγ knockdown on Ovalbumin and cockroach allergens induced AAI

The above findings indicated that RXRγ is essential in lung homeostasis. To determine the involvement of RXRγ in steroid resistance, we downregulated RXRγ in ovalbumin and cockroach allergen induced mice with AAI, with or without steroid treatment. In Ovalbumin set, there were five mice groups: SHAM, OVA, OVA + RXRγ siRNA, OVA + DEX and OVA+ RXRγ siRNA+ DEX as explained in Methods (Model IV, Supplementary Table 1). The allergen induced mice, which otherwise respond to steroids, were found to be poorly responding to the steroids upon downregulation of RXRγ with siRNA. We observed that dexamethasone was not able to alleviate airway hyper-responsiveness in RXRγ siRNA administered OVA mice (Fig. 3A). Also, OVA+ RXRγ siRNA+ DEX mice had increased infiltration of inflammatory cells, mean total inflammation score, and mucous metaplasia compared to OVA+DEX mice (Fig. 3B-C, Fig. S3A). Similarly, dexamethasone reduced OVA specific IgE in OVA mice whereas it did not reduce the same in RXRγ siRNA administered OVA mice (Fig. S3B).

**Figure 3.**
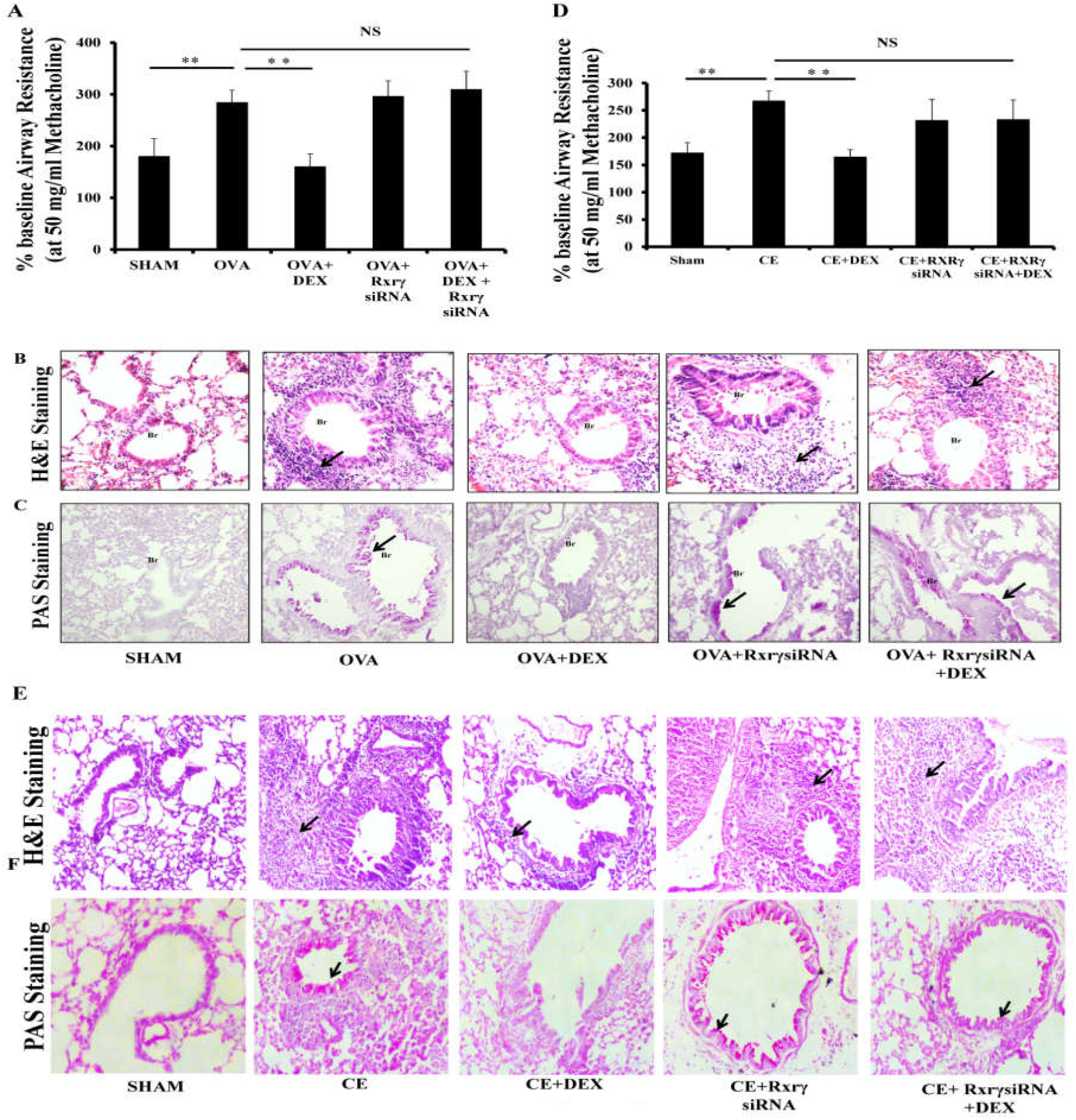
RXRγ Knockdown in allergen induced mice leads to steroid insensitive asthma features. Measurements of percentage baseline airway resistance in response to 50 mg/ml of Methacholine assuming saline aerosol-derived values as baseline in OVA (**A**) and cockroach allergen (**D**) induced mice downregulated with RXRγ by siRNA. **p <0.01 (unpaired t test), NS not significant (unpaired t test). Representative photomicrographs of lung tissue sections stained with H and E (**B & E**) and Periodic-Acid Schiff’s (PAS) (**C & F**) stainings. Black arrows indicate the inflammation in bronchovascular regions in H ad E stained sections and goblet cell metaplasia in PAS stained sections. N=5-6 mice per each group.

Next, we wanted to determine whether RXRγ knockdown could lead to steroid insensitivity in cockroach extract allergen, human relevant allergen, exposed mice with the features of AAI. There were five mice groups: SHAM, CE, CE + RXRγ siRNA, CE + DEX, and CE+ RXRγ siRNA+ DEX as explained in Methods and Supplementary Table 1 (Model V). While DEX treatment reduced airway inflammation, goblet cell metaplasia, total inflammation score and airway hyper-responsiveness in CE mice, it did not in RXRγ siRNA administered CE mice (Fig. 3D-F, Fig. S3C).

We next determined the effects of RXRγ knockdown in allergic mice on BAL fluid cell count and inflammatory cytokines in lung. Dexamethasone treatment reduced both neutrophil and eosinophil percentages in OVA mice whereas it could not reduce them in RXRγ siRNA administered OVA mice (Fig. 4A). However, there was a mild reduction in eosinophils in OVA+ RXRγ siRNA+ DEX mice (Fig. 4A). Similar to OVA model, in CE induced mice, DEX could reduce both neutrophil and eosinophil counts in CE mice. However, it could not reduce neutrophil in RXRγ siRNA administered CE mice (Fig. 4B). However, DEX treatment to RXRγ siRNA administered CE mice was able to reduce eosinophil significantly as compared to CE mice (Fig. 4B). The cytokine analysis revealed that DEX treatment in OVA mice reduced Th2 and Th17 cytokines like IL-13, & IL-17 and KC along with an increase of IFNγ. However, DEX treatment in RXRγ siRNA administered OVA mice could not reduce IL-17 and KC though it reduced IL-13 and IL-5 significantly (Fig. 4C). On the other hand, DEX treatment to CE mice but not RXRγ siRNA administered CE mice decreased IL-17, KC and IL-13 (Fig. 4D). We found an increase in IFNγ levels in CE mice and reduction in CE+ DEX mice with no significant change in CE+ RXRγ siRNA+ DEX mice (Fig. 4D).

**Figure 4.**
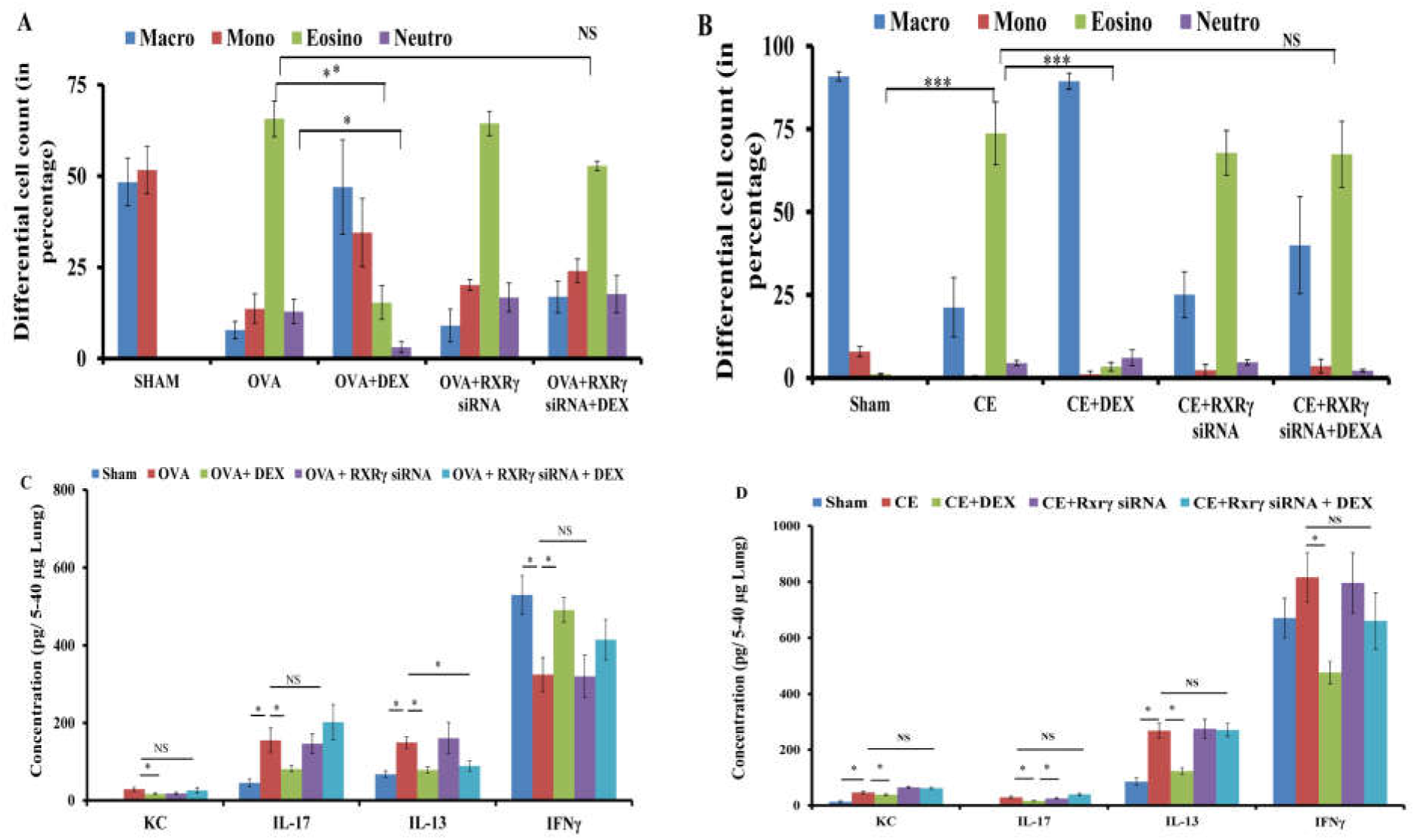
Effects of RXRγ Knockdown in allergen induced mice on BAL cell count and various cytokine in lung. Differential cell count **(A & B), and** the levels of various cytokines in the total lung lysates (**C & D**) of various mice groups (unpaired t test, statistical values have been mentioned above the bars). N=9-10 per each group. Data represents mean ± SEM and this is combined data of two independent experiments; *p <0.05, **p <0.01, ***p <0.001 and NS, not significant

#### RXRγ modulates steroid sensitivity possibly by modulating GRα

To identify the plausible mechanism by which RXRγ might lead to steroid insensitivity, we first checked the effect of RXRγ on GR activation, binding of GR to the synthetic glucocorticoid receptor element (GRE) in nuclear extract of Beas-2B cells pretreated with dexamethasone. We found that the downregulation of RXRγ reduced GR activation and it was not restored with dexamethasone treatment (Fig. 5A). To identify if RXRγ mediated loss of GR activity was due to loss of GRα transcripts, we measured the protein as well as transcript levels of GRα in the Beas-2B cells and observed a drastic decrease in the expression of GRα upon RXRγ downregulation (Fig. 5B-C, Fig. S3D-F). Similarly, GRα was found to be reduced in human asthmatic lungs whereas we did not see any change in the expression of GRβ (Fig. 5D-E). The details of patients and indications of lung resections have been mentioned in Supplementary Table 2.

**Figure 5.**
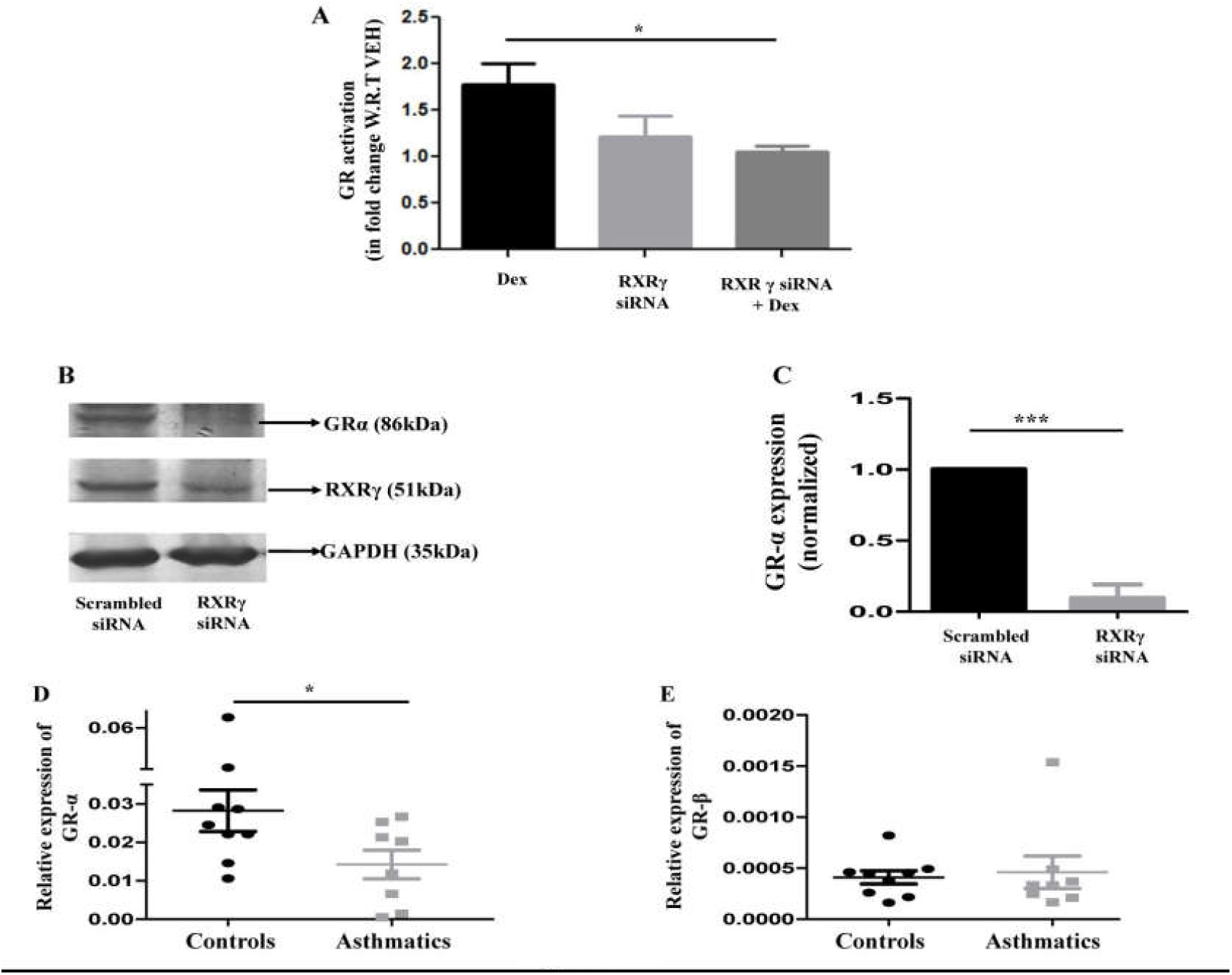
RXRγ knock downed bronchial epithelia had shown the reduction in the glucocorticoid receptor-α expression. **A)** GR activity in Beas-2B cells, treated with or without dexamethasone (10^−6^ M) for 48 hrs after RXRγ siRNA transfection (unpaired t test, this is combined data of 3 independent experiments). The groups are normalized with vehicle/SCR control cells. **B)** Immunoblotting of RXRγ and GRα in the RXRγ siRNA (150nM) treated Beas-2B cell lysates. **C)** GRα transcript levels in the RXRγ siRNA treated Beas-2B cells in fold change with respect to housekeeping genes, beta actin and beta 2 microglobulin (unpaired t test, this is combined data of 4 independent experiments). **D-E)** Measurement of GRα and GRβ transcript levels by qPCR in human lung tissues (unpaired t test, n=8-9 per each group). **F)** Schematic representation of human glucocorticoid receptor gene to show the putative binding sites of RXRγ. **G)** Measurement of GRα transcript levels in Beas-2B cells treated with ATRA (pan agonist of RAR) or 9-cis Retinoic acid (9CRA) or docosahexaenoic acid (DHA) (pan agonists of RXR) by qPCR (unpaired t test and this is combined data of 5 independent experiments). Data represents mean ± SE; *p <0.05, ***p < 0.001. SCR: scrambled, DEX: dexamethasone.

Based on the observation of a reduction in the transcript levels of GRα, we hypothesized that RXRγ could be regulating the expression of GR directly or indirectly. Since there are no such reports on cross-talk of these receptors at the gene level; we performed an in–silico analysis by pattern matching using PERL which identified two putative binding sites for RXR on intron 1 (6047-6064) and intron 2 (95126-95142) of human GR gene (Fig. 6A). While AGGTCA(n)5AGGTCA (∼17bp) was present on intron 1, GGGTCA(n)4AGGTCA was present on intron 2. To ascertain that the putative binding sites are indeed occupied by RXRγ, chromatin immunoprecipitation was performed by pulling down RXR-alpha, beta and gamma from the extracts of BEAS-2B cells. It was observed that while all three isoforms were binding to the intron 1, RXRγ occupancy was more significant on intron 1 (Fig. 6B). Also, there was no recruitment of any of the RXR isoforms on intron 2, exon 1 or the negative control (data not shown). This binding of RXRγ on intron 1 was confirmed using the biotin-streptavidin pull down assay. As compared to the scrambled oligo (SCRE), biotinylated double stranded RXRE, when used as bait, precipitated a significant amount of RXRγ protein, from the nuclear extracts of Beas-2B cells (Fig. 6C).

**Figure 6:**
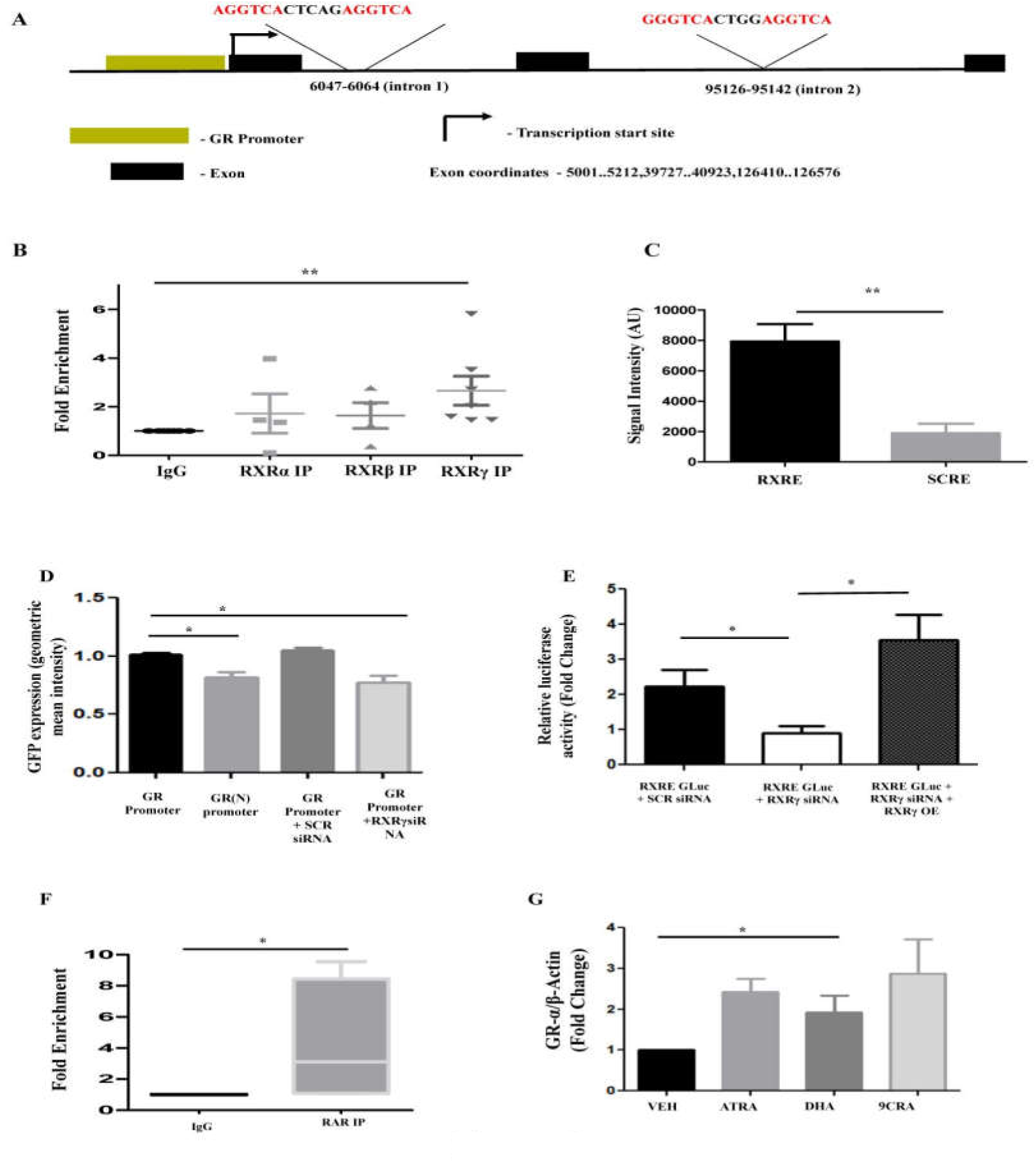
RXRγ regulates glucocorticoid receptor-α. **A)** Schematic representation of human glucocorticoid receptor gene to show the putative binding sites of RXRγ. **B)** Chromatin immuno-precipitation in Beas-2B cells against RXR-alpha, beta and gamma confirms binding of RXR/RAR intron 1 (unpaired t test). **C)** Quantification of oligo-pull down assay showing the occupancy of RXRγ on RXRE of intron 1. **D)** Promoter region of GR Gene including the RXRE cloned upstream of GFP and transfected in Beas-2B cells. GFP expression measured using Flow cytometry. Geometric mean intensity of GFP is plotted for cells transfected with GR promoter alone or with SCR siRNA/RXR-gamma siRNA. Geometric mean for RXRE deleted from intron 1 (GR (N) promoter) was also studied (unpaired t test). **E)** The effect of RXR-gamma knockdown or overexpression on the enhancer was studied by transfecting Beas2b cells with different plasmid constructs for 48hrs as indicated (unpaired t test). **F)** Chromatin immunoprecipitation in Beas-2B cells for pan-RAR on intron 1. Fold enrichment of RAR as compared to IgG antibody (unpaired t test). **G)** Measurement of GRα transcript levels in Beas-2B cells treated with ATRA (pan agonist of RAR) or 9-cis Retinoic acid (9CRA) or docosahexaenoic acid (DHA) (pan agonists of RXR) by qPCR. Data represents mean ± SE; *p <0.05, **p <0.01, ***p < 0.001. SCR: scrambled, GLuc: Gaussea luciferase

Since we identified a binding site for RXRγ on the intron 1 region (will be referred as RXRE from now onwards), we wanted to establish if RXRγ could regulate the expression of GRα. To do so, the entire promoter of GR, exon 1 as well as the intron 1 (∼2.2kb) of GR gene was cloned upstream of an enhanced GFP (EGFP) reporter gene (will be called as GR promoter from now onwards) (Fig. S4B). Transient transfection of the promoter-reporter plasmid (GR promoter) was carried out in Beas-2B cells with or without siRNA mediated downregulation of RXRγ, after which the GFP expression was measured by flow cytometry. It was observed that the mean intensity of EGFP was significantly reduced when RXRγ was downregulated as compared to scrambled oligo or promoter alone (Fig. 6D). To confirm this further, we deleted the 17bp (RXRE) from the intron 1 (GR (N) promoter) (Fig. S4C) and observed a significant reduction in EGFP expression when compared to GR promoter alone (Fig. 6D). This confirms further that the 2.2kb region of GR gene has all the necessary elements to support the transcriptional regulation of GR by RXRγ.

To further confirm that RXRE present in intron 1 conferred transcriptional regulation of GRα, 153 bp, flanking the 17 bp of RXRE was cloned upstream of the HSVTK-promoter with Gaussia luciferase as a reporter gene (RXRE-GLuc) (Fig. S4D-F). Transient co-transfection of RXRE-GLuc and control reporter expressing firefly luciferase was carried out in MCF7 (breast epithelial cells) to normalize the activity of the experimental reporter expression. After normalizing the luciferase activity with the TK-GLuc, it was observed that downregulation of RXRγ reduced the luciferase activity whereas the restoration of RXRγ levels with overexpression retained the luciferase activity (Fig. 6E). Altogether, this indicates that the binding of RXRγ to intron 1 is essential in regulating the expression of GRα and downregulation of RXRγ levels lead to reduced GRα expression.

#### Ligands of RXR-RAR heterodimer positively modulates GRα transcription

Literature suggests that, when the direct repeats of the response element of non-steroidal receptors are spaced by 5 nucleotides, the preferential heterodimerizing partner of RXR is RAR (24). Since the putative binding site on intron 1 of GR gene had a similar configuration, where RXRγ could be heterodimerizing with RAR. To ascertain this, we performed chromatin immunoprecipitation with pan-RAR antibody and observed a significant occupancy on the intron 1 but not intron 2 (Fig. 6F). Further, we wanted to determine the effects of different ligands of RXR-RAR heterodimer on the GRα transcription. There was an increase in the basal transcript levels of GR-α in the BEAS-2B cells, when treated with the naturally occurring ligands of RXR, such as 9-cis Retinoic acid (9-cRA) or docosahexaenoic acid (DHA) or all-trans retinoic acid (ATRA)-the latter having the lowest affinity towards RXR among the three (Fig. 6G). Here, PIAS3 was used as a positive control for the DHA efficiency and we found that there was an increase in the expression of PIAS3 whose promoter is regulated by RXRγ (Fig. S4A).

#### Docosahexaenoic acid (DHA) administration improves steroid sensitivity in murine model of steroid insensitive asthma

DHA or docosahexaenoic acid is considered to be the endogenous ligand for RXR and has been proved to be beneficial in alleviating inflammation. Therefore, we measured the levels of DHA in the sera samples of steroid sensitive and steroid insensitive asthmatic patients. Here, we observed a significant reduction in the levels of DHA in steroid insensitive patients compared to steroid sensitive patients (Fig. 7A). However, there was no reduction in the levels of Retinol, RAR ligand, in steroid insensitive patients as compared to steroid sensitive patients (data not shown). Further, DHA but not ATRA (Model VI, there were 5 mice groups, Fig. S5A) was able to alleviate inflammatory parameters, such as airway hyperresponsiveness, total inflammatory score along with the reduction in both eosinophil and neutrophil counts (Fig. 7B-D, and Fig. S5B). DHA also reduced KC, IL-17, IL-13, and OVA-specific IgE though it could not restore IFNγ significantly (Fig. 7E and Fig. S5C).

**Figure 7.**
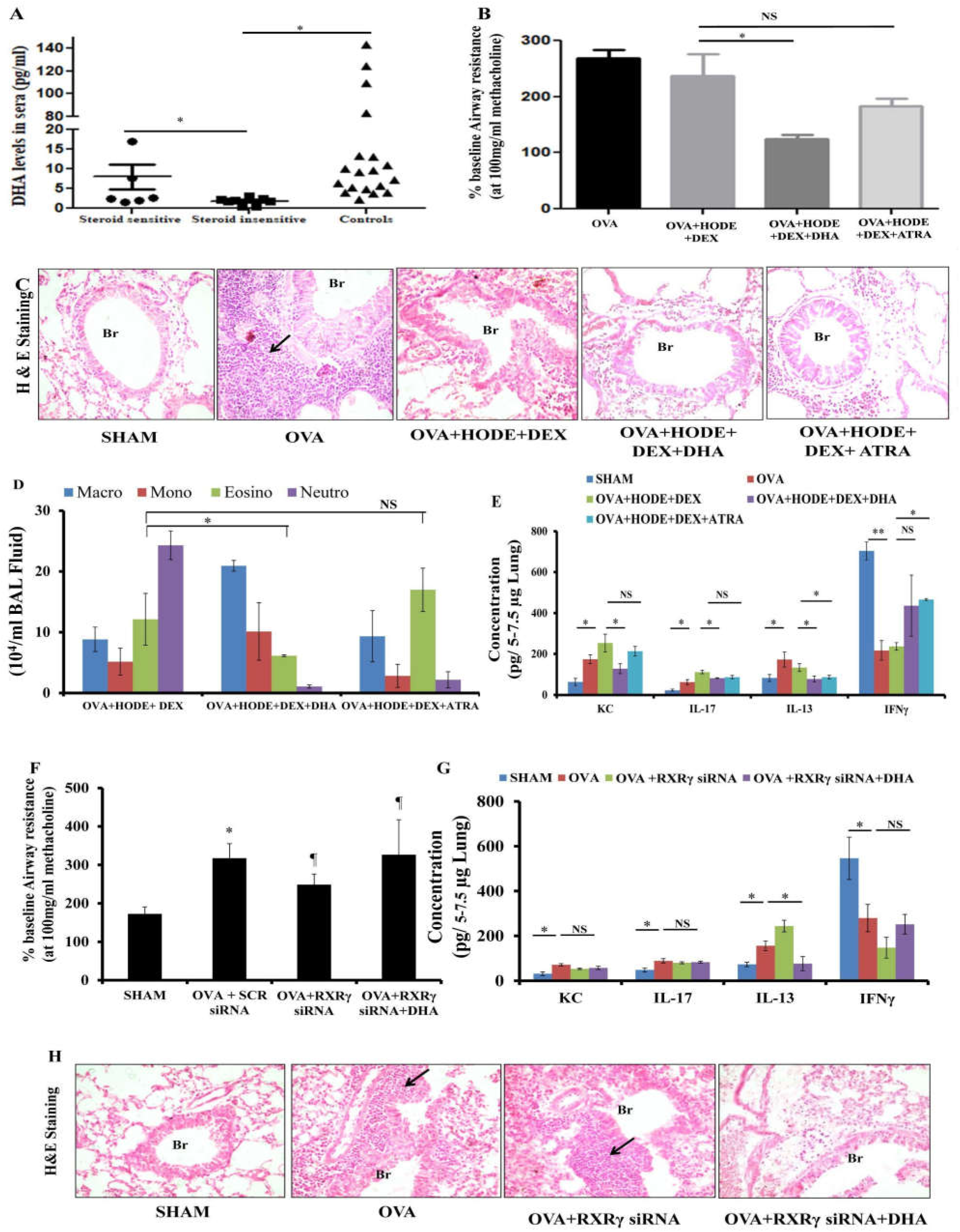
DHA modulates steroid sensitivity. **A)** DHA levels in pg/ml measured in the sera samples of steroid insensitive and steroid sensitive asthmatics (unpaired t test, n=9-25 per each group). **B)** Measurement of percentage baseline airway resistance in response to 100 mg/ml of Methacholine (unpaired t test), **C)** Representative photomicrographs (20 X magnifications) of bronchovascular regions of different groups of mice stained with haematoxylin and eosin (H & E). Black arrows indicate the inflammation in bronchovascular regions. **D)** Absolute cell count in BAL fluid of mice (unpaired t test). **E**) The concentrations of various cytokines like KC, IL-17, IL-13, and IFNγ in the total lung lysates (unpaired t tests, statistical values have been mentioned above the bars). **F)** Measurement of percentage baseline airway resistance in response to 100 mg/ml of Methacholine (unpaired t test). **G**) The concentrations of various cytokines like KC, IL-17, IL-13, IFNγ in the total lung lysates (unpaired t tests, statistical values have been mentioned above the bars). **H)** Representative photomicrographs (20 X magnifications) of bronchovascular regions of different groups of mice stained with haematoxylin and eosin (H & E). Data represents mean ± SEM; n=5-6 per group and shown data is representative of two independent experiments and similar trend was observed in each independent experiment. *p <0.05, **p <0.01. Br : bronchi, NS, Non-significant, * in F panel indicates p <0.05 compared to SHAM group and ¶ indicates Non-significant compared to OVA + SCR siRNA.

**Figure 8.**
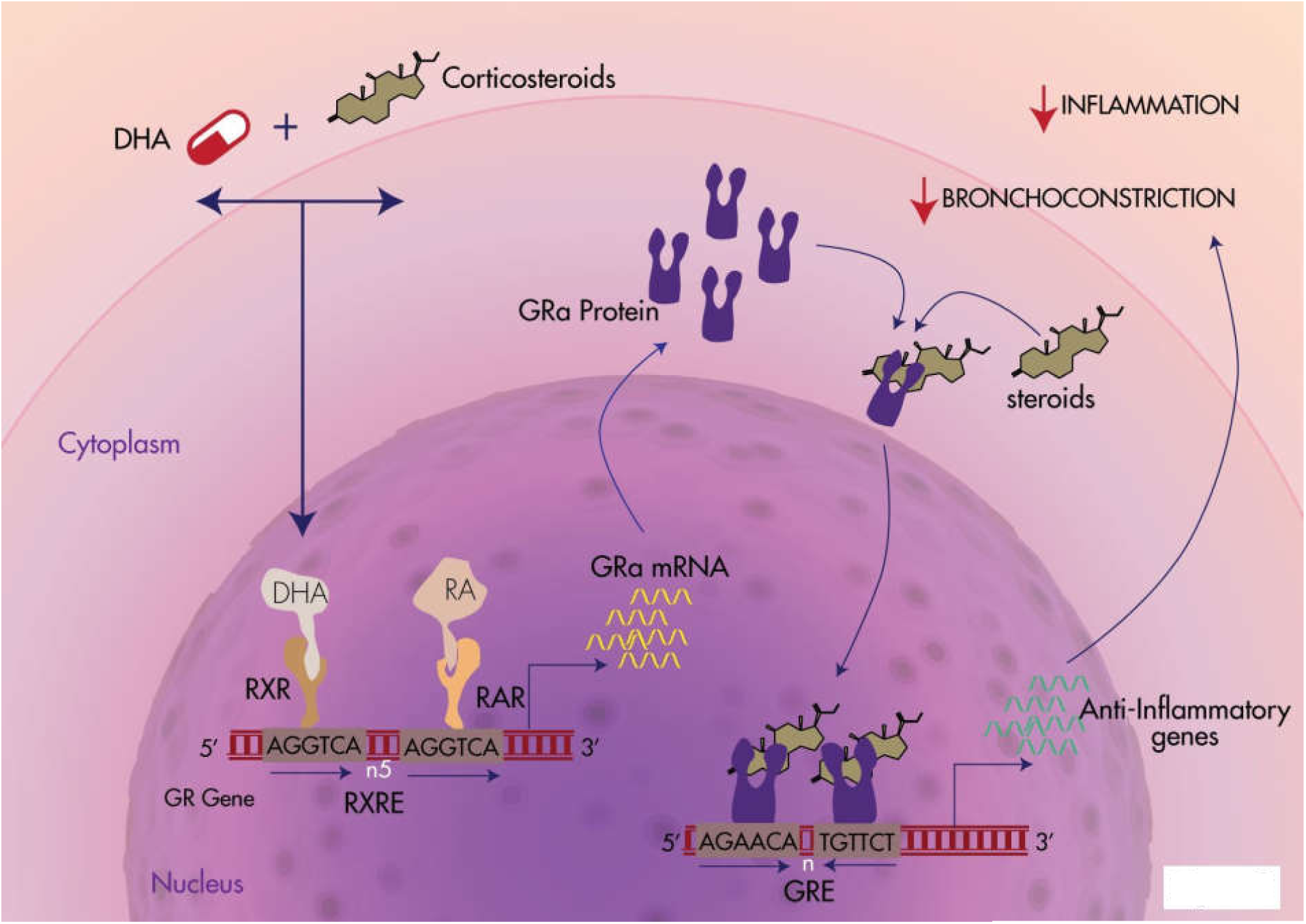
Scheme diagram illustrates the role of RXRγ and its ligand Docosahexaenoic acid (DHA) on modulation of steroid insensitivity in asthma via Glucocorticoid receptor. We have demonstrated the existence of RXR/RAR binding site in glucocorticoid receptor gene. The RXRγ heterodimerizes with RAR to bind to rexinoid receptor element (RXRE) present on intron 1 of glucocorticoid receptor (GR) gene and regulates the expression of GRα. While DHA, a natural ligand of RXR, could increase the expression of GRα, and the resultant increased GR may homodimerize upon corticosteroid treatment and bind to glucocorticoid receptor element (GRE) present on various pro-inflammatory cytokines and anti-inflammatory cytokines genes and reduces the inflammation and also reduction in bronchoconstriction. However, in steroid insensitive condition, when RXRγ levels are low, the regulation on GRα is lost and the expression of GRα is reduced. This further leads to loss of regulation of inflammatory cytokine genes expression and tilting the cellular milieu towards pro-inflammatory and hence, causing steroid insensitivity.

#### Docosahexaenoic acid (DHA) mediated reduction in AHR seems to be RXRγ independent

To study if the effect of anti-inflammatory action of DHA took place via RXRγ, we administered DHA to the Rxrγ knockdown induced steroid insensitive mice (OVA+RXRγ siRNA) (Fig. S5D). There were 4 groups of mice: SHAM, OVA, OVA+RXRγ siRNA and OVA+RXRγ siRNA+DHA (Model VII, Supplementary Table 1). We observed that there was a significant reduction in the total inflammation score and infiltration of inflammatory cells in OVA+RXRγ siRNA+DHA compared to OVA+RXRγ siRNA (Fig. 7H, Fig. S5E). We did not observe any significant change in AHR in OVA+RXRγ siRNA+DHA mice compared to OVA+RXRγ siRNA (Fig. 7F). The cytokine analysis revealed that DHA could not reduce KC, IL-17 and restore IFNγ though it significantly reduced IL-13 and OVA specific IgE (Fig. 7G and Fig. S5F). It is to be noted that Though DHA reduced IL-17 in RXRγ sufficient mice, DHA treatment could not reduce IL-17 in RXRγ knockdown mice. This indicates that DHA mediated airway inflammation is RXRγ independent whereas AHR is RXRγ dependent.

## Discussion

The current study explored the pathophysiological role of RXRγ in steroid insensitive asthma along with the beneficial effects of the DHA supplementation to improve the steroid sensitivity.

Many previously known mechanisms of steroid resistance discuss the involvement of nuclear proteins and receptors, such as HDAC2, PI3K, NF-κB, AP-1, and GR in steroid resistance (25, 26). To begin with, the functioning of steroids is orchestrated in the nucleus hence we studied the status of 84 nuclear genes including nuclear receptors and co-regulators. We observed that only RXRγ showed the expected trend of an anti-inflammatory gene; upregulated with steroid sensitive and downregulated in steroid insensitive condition with an opposite trend in the expression of p100 subunit of NF-κB. This was in concordance with our previous study where we found that HODE induced steroid insensitivity was partially NF-κB dependent (11). As the nuclear events are dynamic with post-transcriptional and post-translational modifications (25, 27, 28), mere gene expression may not capture the whole picture. Therefore, here, we further dissected the mechanistic pathway to reveal how RXRγ plays a role in steroid sensitivity. In contrast to our findings of the reduced expression of RXRγ in the human asthmatic lungs, higher expression of RXRγ has been reported in severe asthmatic patients who were under chronic steroid treatment (29). This is not surprising as steroid itself induces the expressions of RXR and RAR (13) and indeed we also found the increase in the expression of RXRγ in steroid-treated mice with AAI.

The sufficient expression of RXRγ in the lung, especially in the bronchial epithelia, indicates the possible homeostatic role of RXRγ that was evident by spontaneous AHR along with pulmonary inflammation after RXR gamma knockdown in the naïve mice. At the moment, we do not know the exact reason for Rxrγ siRNA mediated spontaneous airway inflammation even in the absence of other stimuli. However, we believe that Rxrγ might control endogenous glucocorticoid receptor signaling that could control sub-clinical inflammatory events. In any event, this needs to be investigated further. While glucocorticoid receptor knockout is lethal, organ or cell-specific conditional knockout mice are highly susceptible to develop inflammation (30). Upon its knockdown with specific siRNA, there was an increase in IL-17 response and IL-4 and decrease in Th1 response without affecting IL-13. As IL-17 is known to act as a neutrophil chemoattractant, this could be attributable to increased airway neutrophilia in Rxrγ siRNA administered mice. However, the detailed mechanism behind this immune response and cellular sources of IL-17 and other cytokines needs to be investigated. The positive relationship between higher IL-17 response and steroid insensitivity is well known (31). Though the primary source of IL-17 is Th17 cells, other cell types like innate lymphoid cells, neutrophils, B cells, natural killer T cells, γδ T cells, even bronchial epithelial cells were shown to secrete IL-17 family cytokines IL-17, IL-22, etc (31). The higher IL-17 response in lungs recruit neutrophils and these neutrophils are activated by CXCL8 or set of chemokines secreted by airway epithelial cells (31). On the other hand, IL-17 also modulates the airway epithelia to induce airway remodeling that is also resistant to steroids. Thus, the role of epithelia in steroid resistance cannot be underestimated. In this context, the expression of Rxrγ in airway epithelia might have a crucial role with respect to steroid resistance.

Severe asthma is not always associated with neutrophilic phenotype but mixed neutrophilic-eosinophilic airway inflammation combined with Th17/Th2 activation has also been reported (31). In this context, we found increased levels of IL-17 family cytokines and IL-4 though IL-13 was not increased in Rxrγ siRNA administered mice. The downregulation of IFNγ in Rxrγ siRNA administered mice may not be surprising in the context of IFNγ attenuating property of IL-17 in airway epithelia (32). Further, RXRγ is also expressed in the other cell types like sub-epithelial mesenchymal cells. Though we have focused on human epithelial cells in this study, we cannot rule out the involvement of non-epithelial cells in HODE or RXRγ siRNA mediated features.

With our in-vitro studies suggesting the regulation of GR by RXRγ, the human asthmatic lungs showing the decreased expression of GRα along with decreased RXRγ; indicate crosstalk between rexinoid and steroid receptors. Though the effects of steroids on the retinoid signaling pathway is well known, the effects of rexinoids on GR and its signaling is not studied. However, ligated RARα/RXR is known to interact with ligated GRα and this interaction further enhances the transcriptional activity of glucocorticoid receptor (GR) regulated genes (12). While this study focused on the protein-protein interaction of retinoic receptors and GR, and GR mediated transcription, it did not discuss the direct transcriptional regulation of GR by the retinoic receptors. In contrast, we not only found the RAR/ RXR binding site in intron 1 of the GR gene but also the functional relevance of this binding site to regulate GR expression in human epithelial cells. While glucocorticoid receptor gene has no distinct TATA or CAAR elements, translation begins only from the 2nd exon. There are multiple promoters on exon 1 and it has been suggested that these multiple promoters in the 2^nd^ exon may regulate the tissue specific expression of GR (28, 33).

Based on our observations in the steroid insensitive murine model, where downregulation of RXRγ led to steroid insensitive like features, and mechanistic studies in bronchial epithelial cells, we suggest that downregulation of the RXRγ could lead to loss of transcriptional regulation of GRα and hence steroid resistance. Meanwhile, similar to the human GR gene, we do observe a putative binding site for RXR/RAR on the mouse GR gene in its first intron (5317-5334bp). Hence, we hypothesize that downregulation of RXRγ might induce steroid resistance via reducing GRα in mice as well. However, it would be essential to study it further as murine segmentation of GRα and GRβ is not as well characterized as in humans (34).

Studies show a relationship between decreased intake of fish oil in the diet with increased risk of asthma (35, 36). The metabolites of DHA and eicosapentaenoic acid (EPA), omega-3 fatty acids such as resolvins and protectins are known to counter regulate eosinophilic airway inflammation. It is always believed that the beneficial effects of DHA are attributable to its balancing of ω3/ω6 ratio. In any event, the beneficial role of DHA in steroid insensitive asthma is novel. In our study, we have used DHA over 9-cis-retinoic acid as a ligand for RXR, as it is more abundant in the human body and it is also accepted as a safe nutritional supplement (15, 37). Though DHA was able to restore steroid sensitivity in HODE-treated steroid insensitive mouse, ATRA could not. While DHA caused the increase in the expression of GR alpha in human bronchial epithelia, DHA supplementation in mice with steroid insensitive condition improved the sensitivity of steroid. Interestingly, asthmatic patients also had shown the reduction in the levels of DHA in sera. Thus, DHA supplementation may be useful in improving steroid insensitive asthmatic condition. Since DHA is a ω-3 fatty acid, it may mediate its anti-inflammatory action through other pathways independent of RXRγ, such as balancing ω-6/ω-3 ratio. Hence, to detect if DHA regulates steroid sensitivity via RXRγ specifically; we treated RXRγ knockdown-OVA mice with DHA and observed a reduction in RXRγ mediated airway inflammation by DHA without affecting airway hyper-responsiveness.

These results indicate that DHA mediated improvement in steroid sensitivity is partially dependent on RXRγ. Overall, DHA mediated reduction in AHR in HODE steroid insensitive mice might be RXRγ dependent whereas DHA mediated reduction in airway inflammation seems to be RXRγ independent. The levels of KC, IL-17, IFNγ were also not significantly reduced similar to AHR in RXRγ siRNA administered DHA treated asthmatic mice. This RXRγ independent reduction of airway inflammation by DHA may be attributable to stabilizing the omega3/omega 6 imbalance. In any case, drawing a conclusion would be difficult since we have not checked if DHA balances the ω-6/ω-3 ratio, for which we would first need to the find out the exact ratio of the ω-6/ω-3 ratio which is responsible for steroid insensitive features. So, this needs further investigations. Nevertheless, it is implied that DHA can improve steroid sensitivity. Interestingly, it has been demonstrated recently that n3 long chain polyunsaturated fatty acids supplementation to pregnant mothers was associated with reduction in occurrence of persistent asthma symptoms (38, 39).

Thus, DHA might be one of the nutritional steroid sensitizing agents and the present study could pave the way for use of DHA supplementation along with steroids to improve steroid sensitivity in steroid insensitive/resistant asthmatic patients and further help in reducing the need of high dose steroid treatment.

## Abbreviations used

AAI: Allergic airway inflammation
AHR: Airway hyper-responsiveness
ATRA: All-trans retinoic acid
AU: Arbitrary units
BAL: Bronchoalveolar lavage
BEAS-2B: Human bronchial epithelial cells
CE: Cockroach allergen extract
ChIP: Chromatin Immunoprecipitation
9CRA: 9-cis retinoic acid
DEX: Dexamethasone
DHA: Docosahexaenoic acid
DMSO: Dimethyl sulfoxide
ELISA: Enzyme-linked immunosorbent assay
GAPDH: Glyceraldehyde 3-phosphate dehydrogenase
GLuc: Gaussia luciferase
GRα: Glucocorticoid receptor alpha
GRE: Glucocorticoid receptor element
H & E: Haematoxylin and eosin
IFNγ: Interferon gamma
IgE: Immunoglobulin
E IL: Interleukin
KC: Keratinocyte chemoattractant
15-LOX: 15-lipoxygenase
MCF-7: Human breast adenocarcinoma cell line
MCP1-α: Monocyte chemoattractant protein1-α
NF-κB: Nuclear Factor kappa-light-chain-enhancer of activated B cells
O.E: Overexpression
OVA: Ovalbumin
p38-MAPK: p38 mitogen-activated protein kinase
PBS: Phosphate buffered saline
RAR: Retinoic acid receptor
PDTC: Pyrrolidinedithiocarbamate
RXRE: Rexinoid receptor element
RXRγ: Retinoid-x-receptor gamma
13-S-HODE: 13*-*hydroxyoctadecadienoic acid
SEM: Standard error mean
siRNA: Small interfering RNA
STAT-6: Signal transducer and activator of transcription 6
Th: T helper
VEH: Vehicle

## Acknowledgments

This work was supported by the projects BSC 0116, MLP 5502, MLP 126 (CSIR), GAP 0118 (Department of Biotechnology) and GAP0084 (Department of Science & Technology) at Institute of Genomics & Integrative Biology and Indian Institute of Chemical Biology, Council of Scientific and Industrial Research, Govt. of India. LP, APJ, DJ, and BKD acknowledge Academy of Scientific and Innovative Research (AcSIR) New Delhi, for their PhD registrations. SC acknowledges Senior Research Fellowship from the Wellcome Trust/DBT India Alliance [grant number 500127/Z/09/Z (2011-17)].

## Author contributions

L.P., and U.M. conceived the idea. L.P., A. P. G., A. J., D. S., B. K. D., R. R., A. R., J. D., S. S., M. K. Y., D. T. S., and U.M. performed the experiments. Y. S. P., P. A. M., and M. V. S. recruited patients for clinical part of the study. L.P., S. C., M. V. S., P. A. M., Y. S. P., S. C., A. A., B. G., U. M. wrote the manuscript.

