## Supplementary Data for "Docosahexaenoic acid (DHA), a nutritional supplement, modulates steroid insensitivity in asthma"

#### Materials and Methods

##### Quantitative Real-Time PCR array

Nuclear receptor array, $RT^2$ Profiler PCR Array, (Qiagen) was performed according to manufacturer's protocol in the total lung mRNA of HODE-treated steroid resistant mice. To do so, we performed a nuclear receptor PCR array which comprised of 86 genes with the total mRNA isolated from the HODE induced steroid resistant mouse lungs. We normalized the gene expression observed in groups OVA, OVA+HODE, OVA+HODE+DEX and OVA+DEX by OVA alone, to nullify the effect observed due to OVA.

### Supplementary Figures with legends

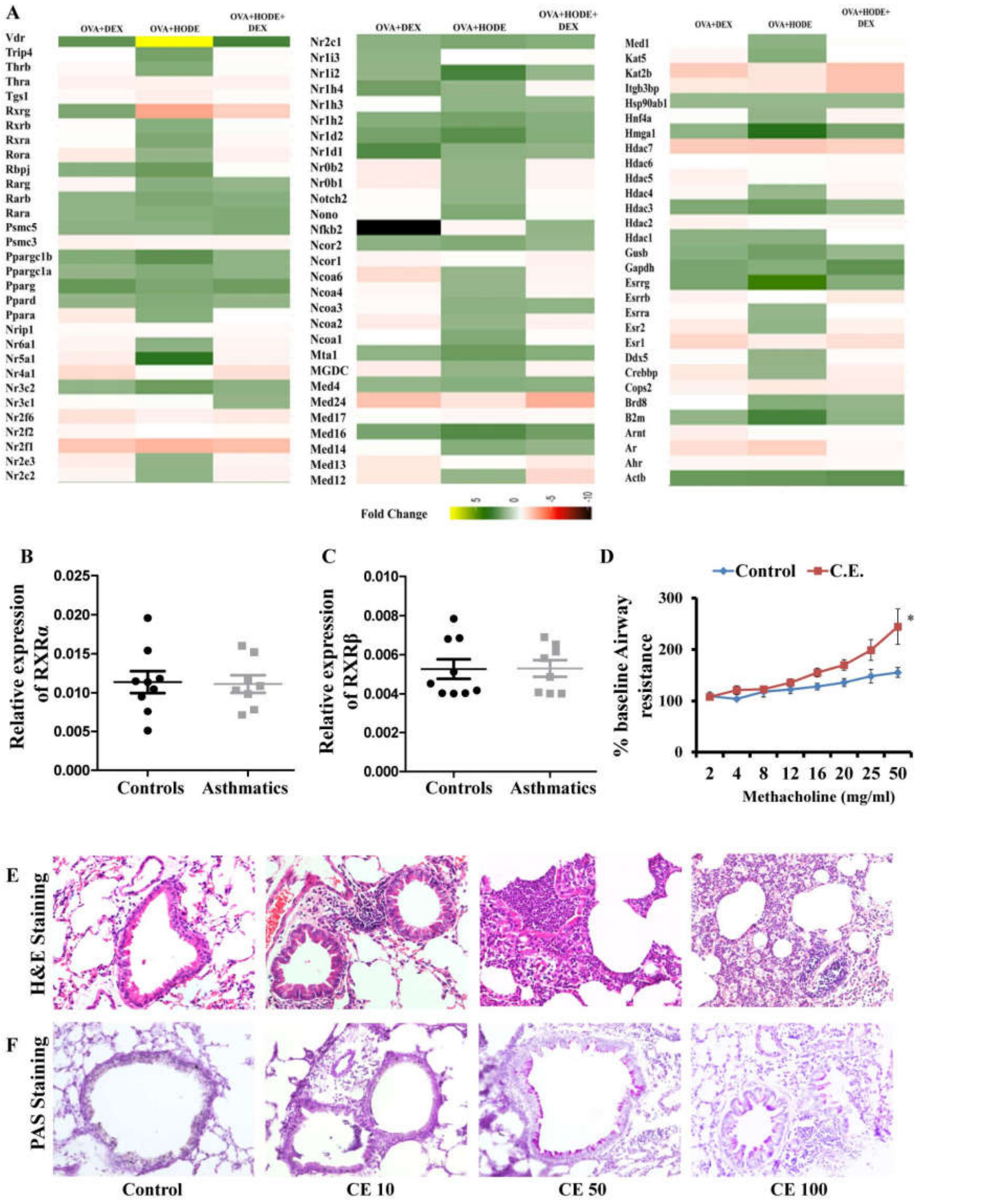


**Figure S1. Heat map of nuclear receptor array in HODE-treated steroid resistant mice lungs and cockroach allergen extract induced allergic airway inflammation. A**) The quantitative Real-Time PCR was performed using nuclear receptor array in the total lung mRNA of HODE-treated steroid resistant mice. We normalized the gene expression observed in groups OVA, OVA+HODE, OVA+HODE+DEX and OVA+DEX by OVA alone, to nullify the effect observed due to OVA. Fold regulation of various groups for the genes shown when compared to OVA group. Transcript levels of RXRα (**B**) and RXRβ (**C**) measured by real time and plotted in fold change with respect to β-actin in total mRNA of human asthmatic and control lungs and unpaired t test was applied in both cases. n=8-9 per each group. The human relevant allergen, cockroach allergen extract (CE), was instilled intranasally at different concentrations like 10μg, 50μg and 100μg to develop the features of allergic airway inflammation in 6-8 weeks old female BALB/c mice. **D**) Measurement of airway resistance in response to increasing concentrations of methacholine as the percent baseline airway resistance assuming saline aerosol-derived values as baseline (n= 5-6 each group). After the end of the protocol, lungs were harvested, processed to get sections and stained with H and E (**E**) and Periodic Acid-Schiff's (**F**) stainings. From these pilot experiments, we had selected 50μg dose for the measurement of AHR. * represents $p < 0.05$ between control and CE groups.

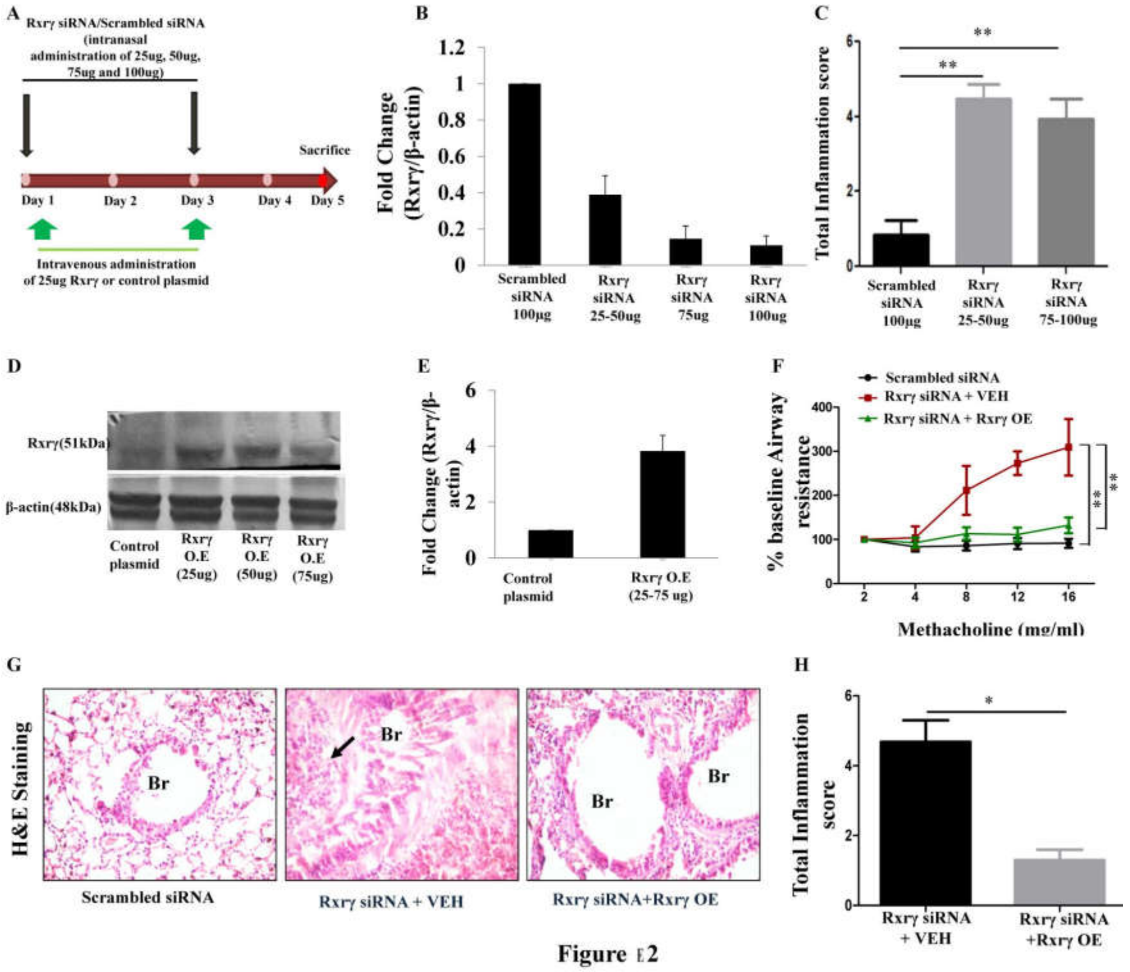


Figure E2

**Fig. S2. Knockdown and overexpression of RXRγ in naïve mice. A)** Schematic representation of experimental design/protocol as described in material and methods (Model III). **B)** Transcript levels of RXRγ after knocking down in mice with different concentrations of siRNA. **C**) Total inflammation score (unpaired t test) of mice groups in naïve mice downregulated with RXRγ by siRNA. The protein (**D**) levels of RXRγ and real time measurement of RXRγ transcripts (**E**) in naïve mice that have been administered with RXRγ overexpression plasmid. Data represents mean ± SE; n= 5-6 each group). **F)** Measurement of airway resistance in response to increasing concentrations of methacholine as the percent baseline airway resistance assuming saline aerosol-derived values as baseline in naïve mice downregulated with RXRγ by siRNA and replenished with RXRγ by overexpression (unpaired t tests between Scrambled siRNA versus RXRγ siRNA + VEH or RXRγ siRNA + VEH versus RXRγ siRNA + RXRγ OE). For statistical calculations, higher dose Methacholine dose (16mg/ml) values were considered. **G)** Representative photomicrographs (20 X magnifications) of bronchovascular regions of different groups of mice stained with haematoxylin and eosin (H & E). **H)** Total inflammation score calculated from H & E stained lung sections (unpaired t test). Data represents mean ± SEM, n=5-10 per group and shown data is representative two independent experiments and similar trend was observed in each independent experiment. Br: Bronchi, VEH : Vehicle, control plasmid that was dissolved in RNAse free water.

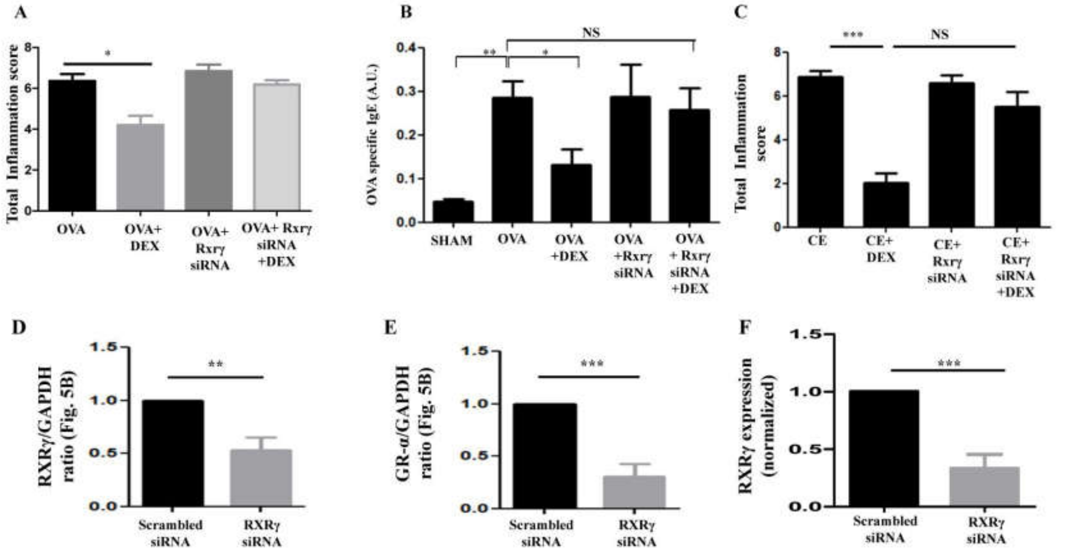


**Fig. S3. Effects of RXRγ siRNA on airway inflammation in allergen induced mice and expression of glucocorticoid receptor-α in Beas2B cells. A)** Total inflammation score (unpaired t test) in OVA induced allergic mice. **B**) OVA specific IgE of various mice groups (unpaired t test, statistical values have been mentioned above the bars). N=9-10 per each group. Data represents mean ± SEM and this is combined data of two independent experiments. **C**) Total inflammation score (unpaired t test) in CE induced allergic mice. Quantification of RXRγ (**D**) and GRα (**E**) in the immunoblot of RXRγ siRNA (150nM) treated Beas-2B total cell lysates using Image J from the figure 5B. **F)** RXRγ transcript levels in the RXRγ siRNA treated Beas-2B cells. *p <0.05, **p <0.01, ***p <0.001 and NS, Non-significant.

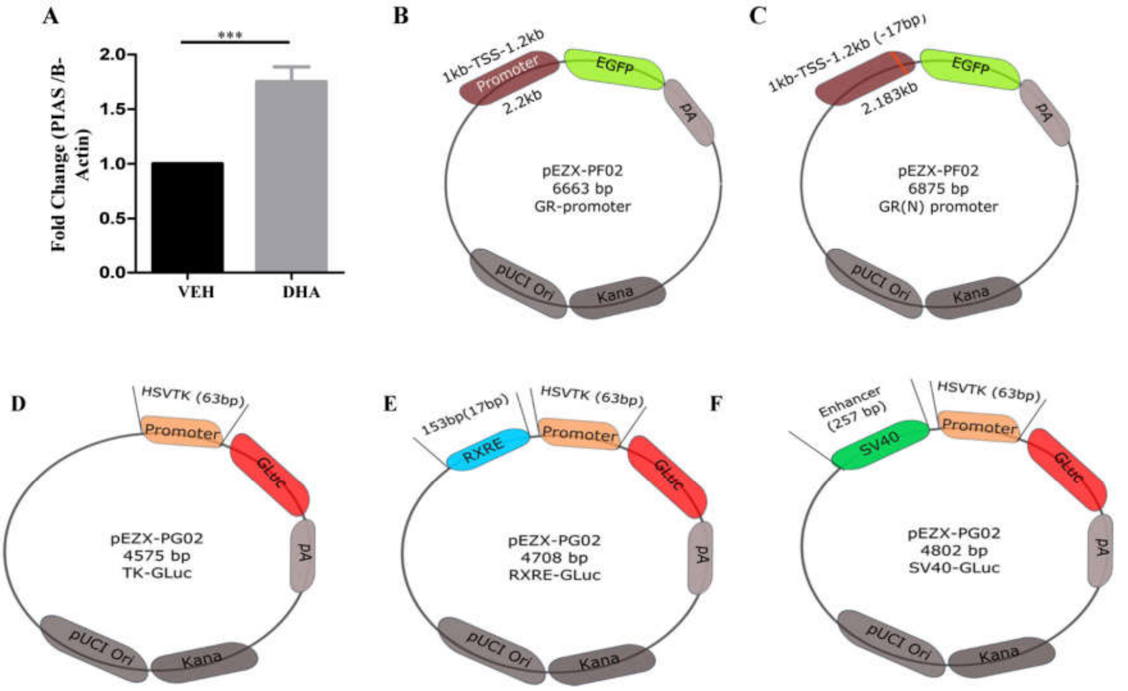


**Fig. S4. RXRγ regulates glucocorticoid receptor-α. A) K)** Transcript levels of PIAS3 upon adding DHA to Beas-2B cells. GR-promoter **(B),** was constructed by cloning 2.2 kb sequence [1kb upstream of transcription start site (TSS) and 1.2 kb downstream of TSS, including intron 1 region with RXR binding element (RXRE) from NR3C1 (GR gene)] upstream of enhanced GFP in pEZX-PF02 vector. GR (N) promoter **(C)** was constructed by cloning the same 2.2kb sequence after deleting RXRE, hence mutant vector in pEZFX-PF02 vector. Luciferase reporter vectors: TK-GLuc, RXRE-GLuc and SV40-GLuc were constructed. TK-GLuc **(D)** was created as negative control with only HSVTK promoter upstream of Gaussia luciferase. RXRE-GLuc **(E)** was constructed by cloning 153 bp sequence flanking RXR binding element (RXRE) from intron1 of NR3C1 upstream of Gaussia luciferase in pEZX-PG02 vector followed by HSVTK promoter (~63bp). SV40-GLuc **(F)** was constructed by cloning 257 bp of SV40 enhancer element upstream of Gaussia luciferase in pEZX-PG02 vector followed by HSVTK promoter, acting as a positive control for the system. Data represents mean ± SE; ***$p < 0.001$. GLuc: Gaussia luciferase.

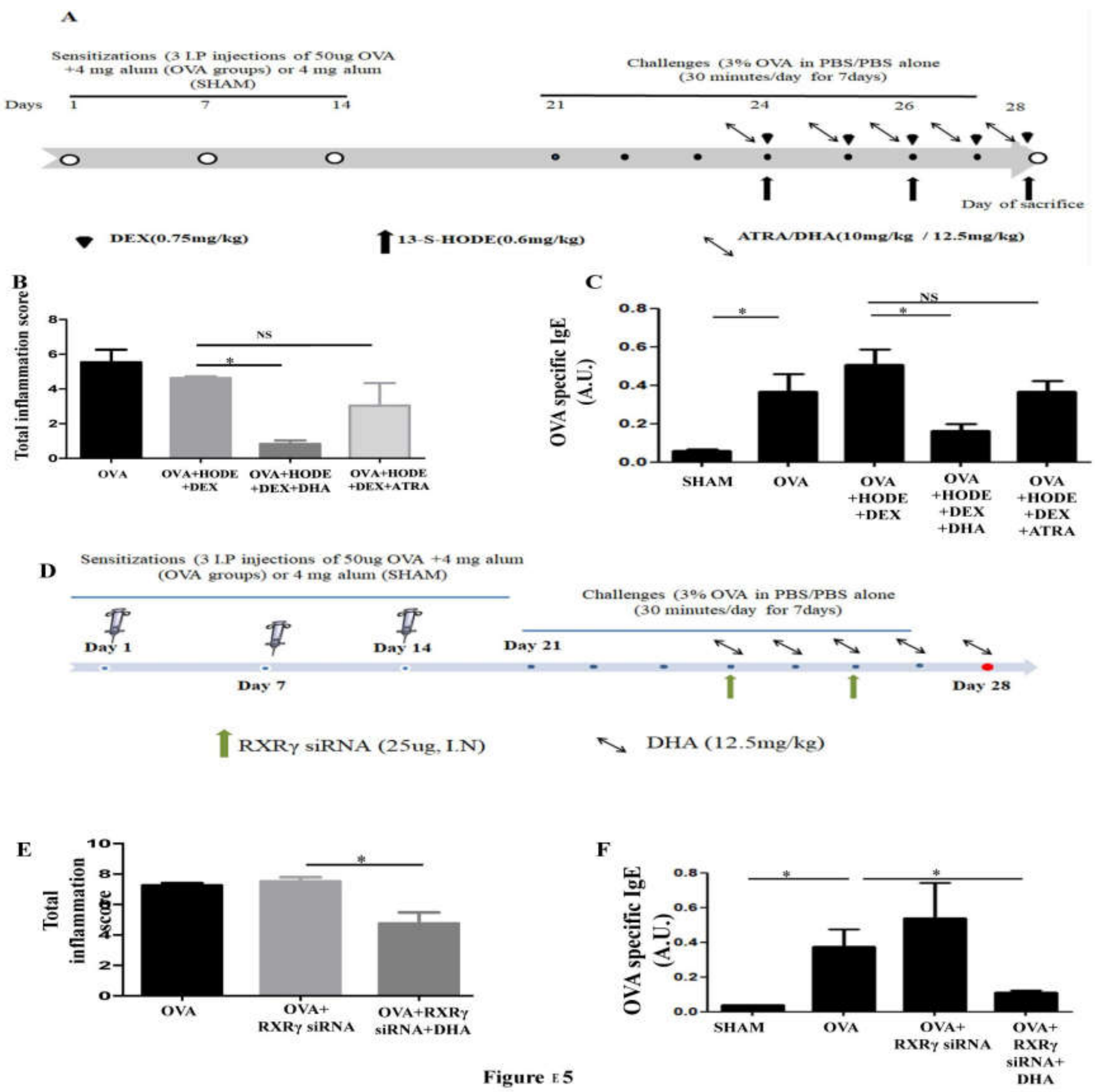


Fig.

**Fig. S5. Effects of DHA airway inflammation in steroid insensitive and RXR knockdown allergic mice. A)** Schematic representation of experimental protocol to determine the effects of DHA in HODE induced steroid insensitive asthmatic mice (Model VI). **B)** Total inflammation score measured in lung sections (unpaired t test). **C**) OVA specific IgE in sera (unpaired t tests, statistical values have been mentioned above the bars). **D)** Schematic representation of experimental protocol to determine the effects of DHA in RXRγ knockdown induced steroid insensitive asthmatic mice (Model VII). **E)** Total inflammation score of the mice groups (unpaired t test). **F**) OVA specific IgE in sera (unpaired t tests, statistical values have been mentioned above the bars). Data represents mean ± SE; *p <0.05 and NS, Non-significant.

#### Supplementary Tables.

| Sr. No | Model | Groups | Group details |
|---|---|---|---|
| I | HODE induced steroid insensitive allergic mice model | A) SHAM<br>B) OVA<br>C) OVA+DEX<br>D) OVA+ HODE<br>E) OVA+HODE+DEX | A) PBS sensitized, PBS challenged and treated with vehicle (50 % ethanol)<br>B) OVA (grade V chicken egg ovalbumin), sensitized, OVA challenged and treated with vehicle<br>C) OVA sensitized, OVA challenged and treated dexamethasone (0.75mg/kg) orally<br>D) OVA sensitized, OVA challenged and intranasal HODE (0.6mg/kg) administered and treated with vehicle<br>E) OVA sensitized, OVA challenged and intranasal HODE (0.6mg/kg) administered and treated with dexamethasone |
| II | Cockroach allergen extract (CE) induced airway inflammation model | A) SHAM<br>B) CE 10<br>C) CE 50<br>D) CE 100 | A) PBS sensitized & challenged<br>B) 10 µg CE sensitized & challenged<br>C) 50 µg CE sensitized & challenged<br>D) 100 µg CE sensitized & challenged |
| III | RXRγ knockdown model in naïve mice | A) Scrambled siRNA<br>B) RXRγ siRNA 25ug<br>C) RXRγ siRNA 75ug<br>D) RXRγ siRNA 100ug | A) Naïve mice intranasally administered with Scrambled siRNA<br>B) Naïve mice intranasally administered with 25ug RXRγ siRNA<br>C) Naïve mice intranasally administered with 75ug RXRγ siRNA<br>D) Naïve mice intranasally administered with 100ug RXRγ siRNA |
| IV | RXRγ knockdown induced steroid insensitive allergic (ovalbumin) mice model | SHAM<br>OVA<br>OVA+DEX<br>OVA+ RXRγ siRNA<br>OVA+ RXRγ siRNA +DEX | A) PBS sensitized, PBS challenged and treated with vehicle (50 % ethanol)<br>B) OVA sensitized, OVA challenged and treated with vehicle<br>C) OVA sensitized, OVA challenged and treated with dexamethasone (0.75mg/kg) orally<br>D) OVA sensitized & challenged, RXRγ siRNA administered and treated with vehicle<br>E) OVA sensitized & challenged, RXRγ siRNA administered and treated with dexamethasone |
| V | RXRγ knockdown induced steroid insensitive allergic (cockroach allergen) mice model | A) SHAM<br>B) CA<br>C) CA +DEX<br>D) CA+ RXRγ siRNA<br>E) CA+ RXRγ siRNA +DEX | A) PBS sensitized, PBS challenged and treated with vehicle (50 % ethanol)<br>B) Cockroach allergen exposed and treated with vehicle<br>C) Cockroach allergen exposed and treated with dexamethasone (0.75mg/kg) orally<br>D) Cockroach allergen exposed, RXRγ siRNA administered and treated with vehicle<br>E) Cockroach allergen exposed, RXRγ siRNA administered and treated with dexamethasone |
| VI | Effects of DHA/ATRA in HODE induced steroid insensitive allergic mice model | A) SHAM<br>B) OVA<br>C) OVA+HODE+DEX<br>D) OVA+HODE+DEX+DHA<br>E) OVA+HODE+DEX+ATRA | A) PBS sensitized, PBS challenged and treated with vehicle (50 % ethanol)<br>B) OVA sensitized, OVA challenged and treated with vehicle<br>C) OVA sensitized, OVA challenged, administered with intranasal HODE and treated with dexamethasone<br>D) HODE treated steroid resistant mice treated with DHA<br>E) HODE treated steroid resistant mice treated with ATRA |
| VII | Effects of DHA in RXRγ siRNA administered allergic mice model | A) SHAM<br>B) OVA<br>C) OVA+ RXRγ siRNA<br>D) OVA+ RXRγ siRNA +DHA | A) PBS sensitized, PBS challenged and treated with vehicle (50 % ethanol)<br>B) OVA sensitized, OVA challenged and treated with vehicle<br>C) OVA sensitized & challenged, and RXRγ siRNA administered<br>D) OVA sensitized & challenged, RXRγ siRNA administered and treated with DHA |

**Supplementary Table 1. Experimental mice models used in this study.**

| Sample codes | Gender | Age | Reason for Surgery | Status | Other pulmonary disease | Smoking History | Co-morbidities |
|---|---|---|---|---|---|---|---|
| **661** | F | 76 | lung nodule | Asthma | None | Past Smoker>20 years | Systemic Hypertension, Hypercholesterolemia, Diabetes |
| **713** | M | 45 | pulmonary nodule | Asthma | None | Current Smoker | Obesity |
| **733** | F | 56 | lung nodule | Asthma | None | Unknown Smoking History | Hypercholesterolemia |
| **752** | F | 68 | lung mass | Asthma | Pulmonary Hypertension | Unknown Smoking History | Systemic Hypertension, Diabetes, Obesity |
| **807** | F | 47 | lung nodule | Asthma | None | Unknown Smoking History | Obesity, Endocrine disease |
| **841** | F | 38 | lung nodule | Asthma | None | Unknown Smoking History | Hypercholesterolemia |
| **848** | f | 24 | lung carcinoma | Asthma | None | Unknown Smoking History | Obesity |
| **850** | F | 41 | metastatic colon cancer | Asthma | None | Non-Smoker | None |
| **354** | M | 54 | lung nodule | Asthma | None | Unknown Smoking History | Obesity |
| **462** | F | 64 | lung nodule | Asthma | None | Non-Smoker | None |
| **615** | F | 73 | carcinoid tumor | Asthma | None | Past Smoker>20 years | Obesity |
| **791** | F | 70 | pulmonary nodule | Normal | None | Non-Smoker | Systemic Hypertension, Obesity |
| **793** | M | 71 | pulmonary nodule | Normal | None | Non-Smoker | None |
| **806** | F | 66 | lung mass | Normal | None | Non-Smoker | Obesity |
| **811** | M | 53 | lung nodule | Normal | None | Non-Smoker | Obesity |
| **845** | F | 57 | pulmonary nodule | Normal | None | Non-Smoker | Heart disease |
| **851** | F | 32 | metastatic colon cancer | Normal | None | Non-Smoker | None |
| **852** | M | 68 | adenocarcinoma | Normal | None | Non-Smoker | Bleeding disorder |
| **853** | F | 68 | carcinoid tumor | Normal | None | Non-Smoker | Systemic Hypertension, Hypercholesterolemia, Heart disease, Obesity |
| **803** | M | 73 | pulmonary nodule | Normal | None | Non-Smoker | Systemic Hypertension, Hypercholesterolemia, Diabetes, Coronary artery disease |
| **822** | F | 59 | leiomyosarcoma | Normal | None | Non-Smoker | Systemic Hypertension, Hypercholesterolemia, Obesity |

**Supplementary Table 2. Brief details of lung samples of human patents used in this study.**

| Sr.No | Gene Symbol | Gene name | OVA+DEX | OVA+HODE | OVA+HODE+DEX |
|---|---|---|---|---|---|
| 1 | Ahr | Aryl-hydrocarbon receptor | -1.2924 | -1.2746 | -1.2226 |
| 2 | Ar | Androgen receptor | -1.879 | -2.1886 | -1.2483 |
| 3 | Arnt | Aryl hydrocarbon receptor nuclear translocator | -1.5052 | -1.007 | -1.1251 |
| 4 | Brd8 | Bromodomain containing 8 | -1.1019 | 1.3195 | 1.021 |
| 5 | Cops2 | COP9 (constitutive photomorphogenic) homolog, subunit 2 | -1.4439 | -1.6472 | -1.5476 |
| 6 | Crebbp | CREB binding protein | -1.6021 | 1.0281 | -1.4641 |
| 7 | Ddx5 | DEAD (Asp-Glu-Ala-Asp) box polypeptide 5 | -1.0425 | 1.0867 | -1.1408 |
| 8 | Esr1 | Estrogen receptor 1 (alpha) | -2.0562 | -1.5052 | -1.8532 |
| 9 | Esr2 | Estrogen receptor 2 (beta) | -1.5692 | 1.1019 | -1.4142 |
| 10 | Esrra | Estrogen related receptor, alpha | -1.1173 | 1.021 | -1.1096 |
| 11 | Esrrb | Estrogen related receptor, beta | -1.3755 | -1.1096 | -1.6133 |
| 12 | Esrrg | Estrogen-related receptor gamma | 1.5369 | 4.3772 | 1.3287 |
| 13 | Hdac1 | Histone deacetylase 1 | 1.2058 | 1.2058 | -1.0867 |
| 14 | Hdac2 | Histone deacetylase 2 | -1.4439 | -1.1647 | -1.2397 |
| 15 | Hdac3 | Histone deacetylase 3 | 1.454 | 1.879 | 1.0867 |
| 16 | Hdac4 | Histone deacetylase 4 | -1.2924 | 1.007 | -1.3379 |
| 17 | Hdac5 | Histone deacetylase 5 | -1.4241 | | -1.1975 |
| 18 | Hdac6 | Histone deacetylase 6 | -1.0792 | -1.0425 | -1.2058 |
| 19 | Hdac7 | Histone deacetylase 7 | -2.3457 | -2.4453 | -2.1886 |
| 20 | Hmga1 | High mobility group AT-hook 1 | 1.1487 | 3.1383 | 1.5801 |
| 21 | Hnf4a | Hepatic nuclear factor 4, alpha | -1.0281 | 1.0644 | -1.366 |
| 22 | Itgb3bp | Integrin beta 3 binding protein (beta3-endonexin) | -1.6586 | -1.7411 | -2.5315 |
| 23 | Kat2b | K(lysine) acetyltransferase 2B | -2.3134 | -1.6935 | -2.5491 |
| 24 | Kat5 | K(lysine) acetyltransferase 5 | -1.2658 | 1.3287 | -1.0792 |
| 25 | Med1 | Mediator complex subunit 1 | -1.0943 | 1.1975 | -1.0644 |
| 26 | Med12 | Mediator of RNA polymerase II transcription, subunit 12 homolog (yeast) | -1.6702 | 1.0281 | -2.042 |
| 27 | Med13 | Mediator complex subunit 13 | -1.6021 | -1.1329 | -1.5692 |
| 28 | Med14 | Mediator complex subunit 14 | -1.0497 | 1.3379 | 1.0497 |
| 29 | Med16 | Mediator complex subunit 16 | 1.5911 | 2.3295 | 1.7291 |
| 30 | Med17 | Mediator complex subunit 17 | -1.0644 | -1.2311 | -1.2142 |
| 31 | Med24 | Mediator complex subunit 24 | -2.5315 | -1.7171 | -2.9897 |
| 32 | Med4 | Mediator of RNA polymerase II transcription, subunit 4 homolog (yeast) | 1.0425 | 1.257 | 1.2226 |
| 33 | Mta1 | Metastasis associated 1 | 1.1173 | 1.8661 | 1.3104 |
| 34 | Ncoa1 | Nuclear receptor coactivator 1 | -1.0943 | 1.4142 | -1.1408 |
| 35 | Ncoa2 | Nuclear receptor coactivator 2 | -1.5263 | 1.057 | -1.3851 |
| 36 | Ncoa3 | Nuclear receptor coactivator 3 | -1.1647 | 1.2058 | 1.1096 |

| | | | | | |
|---|---|---|---|---|---|
| 37 | Ncoa4 | Nuclear receptor coactivator 4 | -1.2483 | 1.0425 | -1.2746 |
| 38 | Ncoa6 | Nuclear receptor coactivator 6 | -1.9053 | 1.021 | -1.3566 |
| 39 | Ncor1 | Nuclear receptor co-repressor 1 | -1.366 | -1.0425 | -1.3947 |
| 40 | Ncor2 | Nuclear receptor co-repressor 2 | 1.2142 | 1.4142 | 1 |
| 41 | Nfkb2 | Nuclear factor of kappa light polypeptide gene enhancer in B-cells 2, p49/p100 | -10.4107 | -1.3104 | 1.0792 |
| 42 | Nono | Non-POU-domain-containing, octamer binding protein | -1.0425 | 1.3947 | -1.0281 |
| 43 | Notch2 | Notch gene homolog 2 (Drosophila) | -1.1329 | 1.1329 | -1.1251 |
| 44 | Nr0b1 | Nuclear receptor subfamily 0, group B, member 1 | -1.5263 | 1.0644 | -1.366 |
| 45 | Nr0b2 | Nuclear receptor subfamily 0, group B, member 2 | -1.5263 | 1.0644 | -1.366 |
| 46 | Nr1d1 | Nuclear receptor subfamily 1, group D, member 1 | 2.3784 | 1.1728 | 1.0718 |
| 47 | Nr1d2 | Nuclear receptor subfamily 1, group D, member 2 | 1.6818 | 2.2191 | 1.3472 |
| 48 | Nr1h2 | Nuclear receptor subfamily 1, group H, member 2 | 1.2924 | 1.6472 | 1.2483 |
| 49 | Nr1h3 | Nuclear receptor subfamily 1, group H, member 3 | -1.0497 | 1.1487 | 1.0353 |
| 50 | Nr1h4 | Nuclear receptor subfamily 1, group H, member 4 | 1.7532 | 1.057 | -1.2834 |
| 51 | Nr1i2 | Nuclear receptor subfamily 1, group I, member 2 | 1.0792 | 2.5669 | 1.021 |
| 52 | Nr1i3 | Nuclear receptor subfamily 1, group I, member 3 | 1.0943 | -1.1096 | -1.181 |
| 53 | Nr2c1 | Nuclear receptor subfamily 2, group C, member 1 | 1.1647 | 1.5052 | 1.2746 |
| 54 | Nr2c2 | Nuclear receptor subfamily 2, group C, member 2 | -1.3472 | 1.1173 | -1.3947 |
| 55 | Nr2e3 | Nuclear receptor subfamily 2, group E, member 3 | -1.5263 | 1.0644 | -1.366 |
| 56 | Nr2f1 | Nuclear receptor subfamily 2, group F, member 1 | -2.4794 | -2.8481 | -2.6759 |
| 57 | Nr2f2 | Nuclear receptor subfamily 2, group F, member 2 | -1.3104 | -1.0792 | -1.1728 |
| 58 | Nr2f6 | Nuclear receptor subfamily 2, group F, member 6 | -1.8025 | -1.3947 | -1.5911 |
| 59 | Nr3c1 | Nuclear receptor subfamily 3, group C, member 1 | -1.1728 | -1.021 | 1.057 |
| 60 | Nr3c2 | Nuclear receptor subfamily 3, group C, member 2 | 1.0943 | 1.8025 | 1.1647 |
| 61 | Nr4a1 | Nuclear receptor subfamily 4, group A, member 1 | -1.9453 | -1.1329 | -1.8532 |
| 62 | Nr5a1 | Nuclear receptor subfamily 5, group A, member 1 | -1.5157 | 3.0951 | -1.366 |
| 63 | Nr6a1 | Nuclear receptor subfamily 6, group A, member 1 | -1.3566 | 1.2058 | -1.3195 |
| 64 | Nrip1 | Nuclear receptor interacting protein 1 | -1.1892 | -1.1892 | -1.2226 |
| 65 | Ppara | Peroxisome proliferator activated receptor alpha | -1.5692 | 1.2924 | -1.1096 |
| 66 | Ppard | Peroxisome proliferator activator receptor delta | 1.057 | 1.3755 | 1.0644 |
| 67 | Pparg | Peroxisome proliferator activated receptor gamma | 1.9588 | 1.5052 | 1.8277 |
| 68 | Ppargc1a | Peroxisome proliferative activated receptor, gamma, coactivator 1 alpha | 1.0718 | 1.3566 | 1.1487 |

| | | | | | |
|---|---|---|---|---|---|
| 69 | Ppargc1b | Peroxisome proliferative activated receptor, gamma, coactivator 1 beta | 1.3755 | 2.1435 | 1.1019 |
| 70 | Psmc3 | Proteasome (prosome, macropain) 26S subunit, ATPase 3 | -1.434 | -1.3379 | -1.3379 |
| 71 | Psmc5 | Protease (prosome, macropain) 26S subunit, ATPase 5 | 1.2226 | 1.2142 | 1.4142 |
| 72 | Rara | Retinoic acid receptor, alpha | 1.1408 | 1.3013 | 1.454 |
| 73 | Rarb | Retinoic acid receptor, beta | 1.1019 | 1.4641 | 1.2397 |
| 74 | Rarg | Retinoic acid receptor, gamma | -1.3566 | 1.2397 | 1.0353 |
| 75 | Rbpj | Recombination signal binding protein for immunoglobulin kappa J region | 1.2483 | 1.9053 | -1.057 |
| 76 | Rora | RAR-related orphan receptor alpha | -1.5911 | 1.0792 | -1.4044 |
| 77 | Rxra | Retinoid X receptor alpha | -1.0718 | 1.4743 | -1.257 |
| 78 | Rxrb | Retinoid X receptor beta | -1.1487 | 1.2483 | -1.021 |
| 79 | Rxrg | Retinoid X receptor gamma | **1.5263** | **-3.1821** | **-2.2658** |
| 80 | Tgs1 | Trimethylguanosine synthase homolog (S. cerevisiae) | -1.2058 | -1.454 | -1.1251 |
| 81 | Thra | Thyroid hormone receptor alpha | -1.3851 | -1.4142 | -1.3851 |
| 82 | Thrb | Thyroid hormone receptor beta | -1.2483 | 1.3566 | -1.007 |
| 83 | Trip4 | Thyroid hormone receptor interactor 4 | -1.1096 | 1.6472 | -1.1408 |
| 84 | Vdr | Vitamin D receptor | 2.1287 | 8.2249 | 2.639 |

**Supplementary Table 3. The profile of differentially expressed genes in Steroid insensitive mice**
